# High-field fMRI at 7 Tesla reveals topographic responses tuned to number in the developing human brain

**DOI:** 10.1101/2025.01.10.632353

**Authors:** Ga-Ram Jeong, Joram Soch, Robert Trampel, Andreas Nieder, Michael A. Skeide

## Abstract

In the adult brain, hemodynamic responses to visually presented numerosities reveal receptive field-like tuning curves in topographically organized maps across association cortices. It is currently unknown whether such tuned topographic responses to numerosities can also be detected in the developing brain. Here we conducted a 7 Tesla fMRI experiment in which we presented a large set of visual dot displays to 11-12-year-old children and adults. We found that developing hemodynamic responses indeed revealed logarithmic Gaussian tuning to quantitative information. Tuning models explained comparable amounts of variance in children and adults. In most subjects, six bilateral cortical maps consistently exhibited topographic responses to numerosities. The present study goes beyond previous work by uncovering a population code for quantity detection in individual developing human brains. Our work lays a foundation for a model-based neuroimaging approach to individual cognitive differences in the context of developmental dyscalculia and mathematical giftedness.

## Main

Looking time experiments suggest that humans are born with a basic ability to discriminate the number of items in two sets without counting^1^. This capacity for approximate numerosity discrimination in newborns is initially limited to a 1:3 ratio (4 versus 12 items), but increases substantially with further age. Six-month-old infants can already discriminate numerosities with a ratio of 1:2 (such as 8 compared to 16 items)^2^. Ten-month-old infants are able to discriminate numerosities with a 2:3 ratio, but not yet a 4:5 ratio^3^. Six-year-old children master a 5:6 ratio and adults even master a ratio of up to 9:10^4^. These group-average age effects on number recognition are accompanied by pronounced individual differences^4^.

In line with the behavioral findings there is electroencephalography evidence for neurophysiological discrimination of 4 versus 12 items in infants as young as 3 months^5^. Moreover, hemodynamic responses of 6-months-old infants undergoing near-infrared spectroscopy suggest that, at this age, the developing brain is able to discriminate between 8 and 16 items^6^. By the age of 3 years at the latest, hemodynamic responses are modulated by a 2:3 ratio including differences between 12 and 16 as well as 16 and 24 items^7^. Finally, functional magnetic resonance imaging data of 12-year-old children indicate neural sensitivity to a 5:7 ratio^8^. Given that these studies employed group-level averaging, individual differences underlying number representation in the developing brain remain poorly understood.

Experiments that are based on the comparison of two sets of items have significantly advanced our understanding of how number information is encoded in the brain. At the same time, these paradigms inherently reduce number recognition to categorical binary classification problems. Detecting adjacent items that are represented in a continuous space, however, is a defining feature of number recognition. In fact, continuous representation of adjacent items is the foundation for understanding core numerical concepts such as magnitude and order (cardinality and ordinality)^9^.

In adults, electrophysiological recordings from single neurons of neurosurgical patients revealed populations with tuned responses to the adjacent numerosities one to five. Specifically, neuronal response profiles revealed peak activity for one preferred numerosity and a gradual decrease of activity the more the number of dots deviated from the preferred value. This pattern followed logarithmic Gaussian tuning curves similar to those known from visual receptive fields^10,11^. Hemodynamic response profiles similar to these electrophysiological response profiles were also found at the whole-brain level by high-field functional magnetic resonance imaging in adults. Remarkably, these tuned responses formed six bilateral topographic maps in which adjacent numerosities are represented in adjacent patches of cortex^12,13^. Whether tuned responses to adjacent numerosities that form topographic maps already emerge in the developing brain and to which degree they differ from the adult brain, is currently unknown.

To tackle these questions, we recorded high-field event-related functional magnetic resonance imaging data at 7 Tesla in 11-12-year-old children and adults. During the experiment participants viewed a large set of visual dot arrays with varying positions but constant luminance in which the numerosities one to five appeared 384 times each, resulting in 1,920 total trials. Following previous electrophysiological and imaging work with a comparably large number of trials, we employed a single-subject design in which 24 individual data sets (12 children and 12 adults) served as replication units and not as measurement units.

We hypothesized that logarithmic Gaussian tuning and topographically organized responses to adjacent numerosities already emerge in the developing brains of 11-12-year-old children. This hypothesis was based on a functional magnetic resonance imaging study demonstrating that averaged hemodynamic responses of 12-year-old children are already sensitive to differences between numerosities with a 5:7 ratio^8^. Our second hypothesis was that tuning would be less precise in children compared to adults, so that tuning models would explain less hemodynamic signal variance in the developing brain. This hypothesis builds on behavioral evidence that even at the age of 14 years, the precision of number discrimination is on average still significantly below adult levels^4^.

In line with our first hypothesis, across children and adults, we found receptive-field-like Gaussian tuning and topographic responses to numerosity distributed over six bilateral cortical maps. Contrary to our second hypothesis, the tuning models explained comparable amounts of hemodynamic variance in children and adults. We observed considerable individual differences with respect to the variance explained by the models and the spatial distribution of the topographic maps. These effects were not related to behavioral differences in attention.

## Results

### Behavioral task

Participants were instructed to press a button whenever visual dot arrays were shown in different color (Fig. 1). Across all participants and runs, the average hit rate for these catch trials was 97.62% in children and 94% in adults (Supplementary Table S1). This near-ceiling performance indicates that participants were continuously paying attention to the stimuli.

**Fig. 1.**
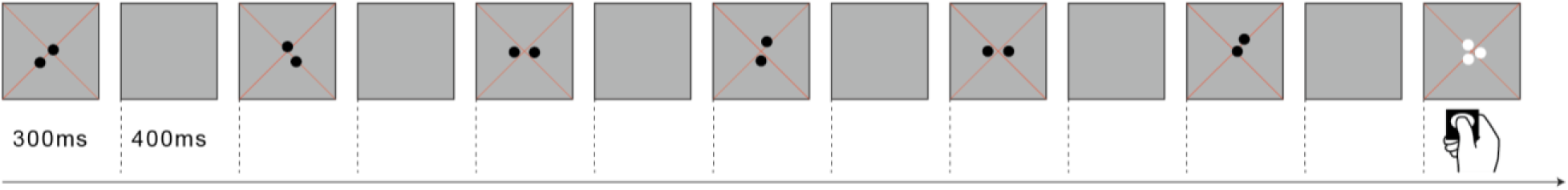
Stimuli and task. Participants viewed six different dot arrays per numerosity that were separately displayed for 300 ms and alternating with 400 ms of gray background. Arrays appeared within a radius of 0.75° (visual angle) of a diagonal cross of thin red fixation lines. Summed surface area was kept constant and dot positions were randomly varied. Participants were instructed to press a button whenever white instead of black dots were shown. These catch trials occurred in 10% of all stimulus events. Numerosities one through five were first presented in ascending order, followed by a block of twenty items, then in descending order, followed by another block of twenty items before this cycle was repeated.

### Logarithmic Gaussian tuning to visual numerosity

Building on previous electrophysiological and magnetic resonance imaging work in humans, we developed population receptive field models describing logarithmic Gaussian functions for numerosity tuning^10-13^. These models were estimated in individual human subjects, separately for each vertex on the cortical surface, after removing possible effects of head motion and nuisance signals (see Methods for details). Our models captured the peak response of a neural population to a preferred numerosity and the numerosity range to which a neuronal population responds (full width at half maximum, FWHM) (Fig. 2a,c,e,g; for replication in other subjects, see Supplementary Fig. S1). Summarizing the hemodynamic activation based on these two parameters, these models explained substantial amounts of the observed signal variance related to regional differences in numerosity tuning (children: maximum cross-validated R² (cvR²) = 0.73, adults: maximum cross-validated R² = 0.72, both p < 0.001) (Fig. 2b,d,f,h; for replication in other subjects, see Supplementary Fig. S1 and S2). The variance explained by the tuning models did not differ significantly between children and adults (t(22) = 0.582, p = 0.566) (Supplementary Table S1).

**Fig. 2.**
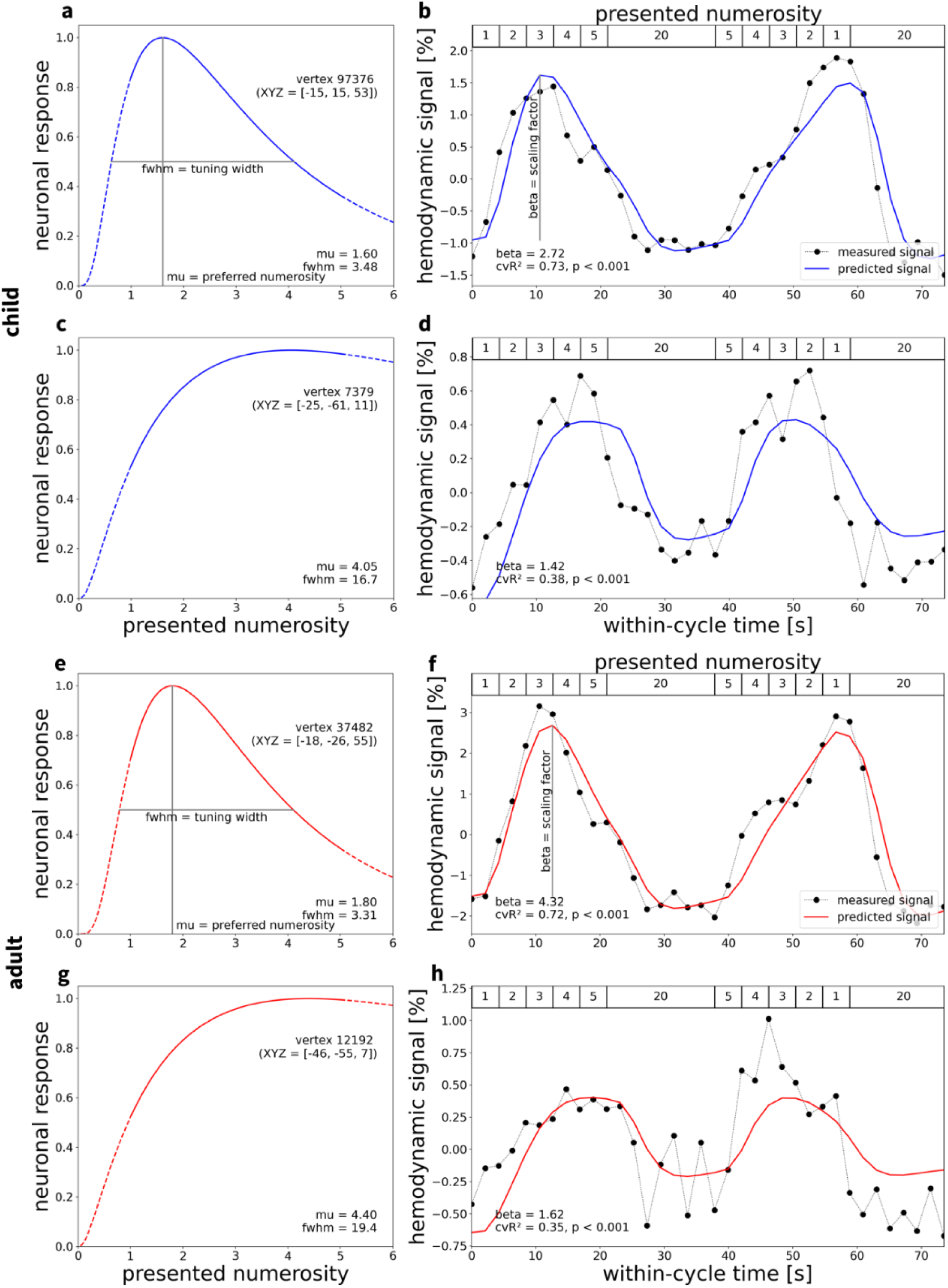
Neural numerosity tuning and hemodynamic activation time course. Results of numerosity population receptive field analyses for **a–d** an example child (top, blue) and **e–h** an example adult (bottom, red). **a–d** depicts vertices in left frontal cortices (**a,b**) and occipital cortices (**c,d**). **e–h** depicts vertices in left parietal (**e,f**) and occipital cortices (**g,h**). For details about anatomical locations, see Supplementary Table S2. Logarithmic Gaussian tuning functions are shown for **a,e** a small-numerosity vertex and **c,g** a larger-numerosity vertex per subject (XYZ = coordinates in FreeSurfer fsnative space). These functions are described by a preferred numerosity (mu) and the full width at half maximum (fwhm). **b,d,f,h**, Combining neuronal tuning models with a hemodynamic forward model generated predicted time courses (solid lines) for numerosity perception. Comparing predicted to measured time courses (dotted lines, averaged across runs and cycles) yielded the cross-validated coefficient of determination (cvR²). This coefficient quantifies the out-of-sample variance explained by the model, trained on averaged odd runs and tested on averaged even runs, and vice versa. For each subject, results are shown for those vertices with the largest cvR² in the small-numerosity range (**a,b,e,f**) and in the large-numerosity range (**c,d,g,h**).

Maximum cvR² across both hemispheres was positively related to the behavioral attentional performance of the adults, but this association was driven by two outlier subjects whose hit rates were about 15% below all others (Bayesian correlation analysis, r_Pearson_ = 0.86, posterior median r = 0.79, 95% CI = [0.47, 0.96], BF_10_ = 115.03, uniform prior ranged [-1,1]). Maximum cvR² was not related to the behavioral attentional performance of the children (Bayesian correlation analysis, r_Pearson_ = 0.41, posterior median r = 0.34, 95% CI = [-0.20, 0.75], BF_10_ = 0.79, uniform prior range [-1,1]).

Although the experimental stimulation only included whole numbers (e.g. not three and a half dots), the numerosity tuning model estimated non-integer numerosities for most vertices (Fig. 2). These results were expected since the large neuronal population sampled by each vertex combines nerve cells exhibiting different preferred numerosities. Accordingly, the preferred numerosity of each vertex roughly corresponds to the expected value across all cells.

### Spatial distribution of hemodynamic responses to visual numerosity

In each hemisphere, we identified six visual regions in which the numerosity tuning models explained measured signal variance that was significantly different from zero (p < 0.05, Bonferroni-corrected for the number of vertices; see Methods for details) (Fig. 3a,b; for replication in other subjects, see Supplementary Fig. S3). In accordance with the literature^12^, we denote these numerosity-selective maps using abbreviations starting with “N”. Tuning was evident in temporo-occipital (NTO), parieto-occipital (NPO), posterior parietal (NPC1), dorsal anterior parietal (NPC2), ventral anterior parietal (NPC3) and superior frontal (NF) cortices (Fig. 3c,d). These numerosity-specific regions were found in a majority of subjects (Fig. 4a,b), but revealed considerable individual differences (Fig. 3c,d). Within numerosity-selective regions, numerosities preferred by vertices, i.e. stimulus intensities for which vertices revealed their peak response, formed topographic maps (Fig. 3e,f; for replication in other subjects, see Supplementary Fig. S4).

**Fig. 3.**
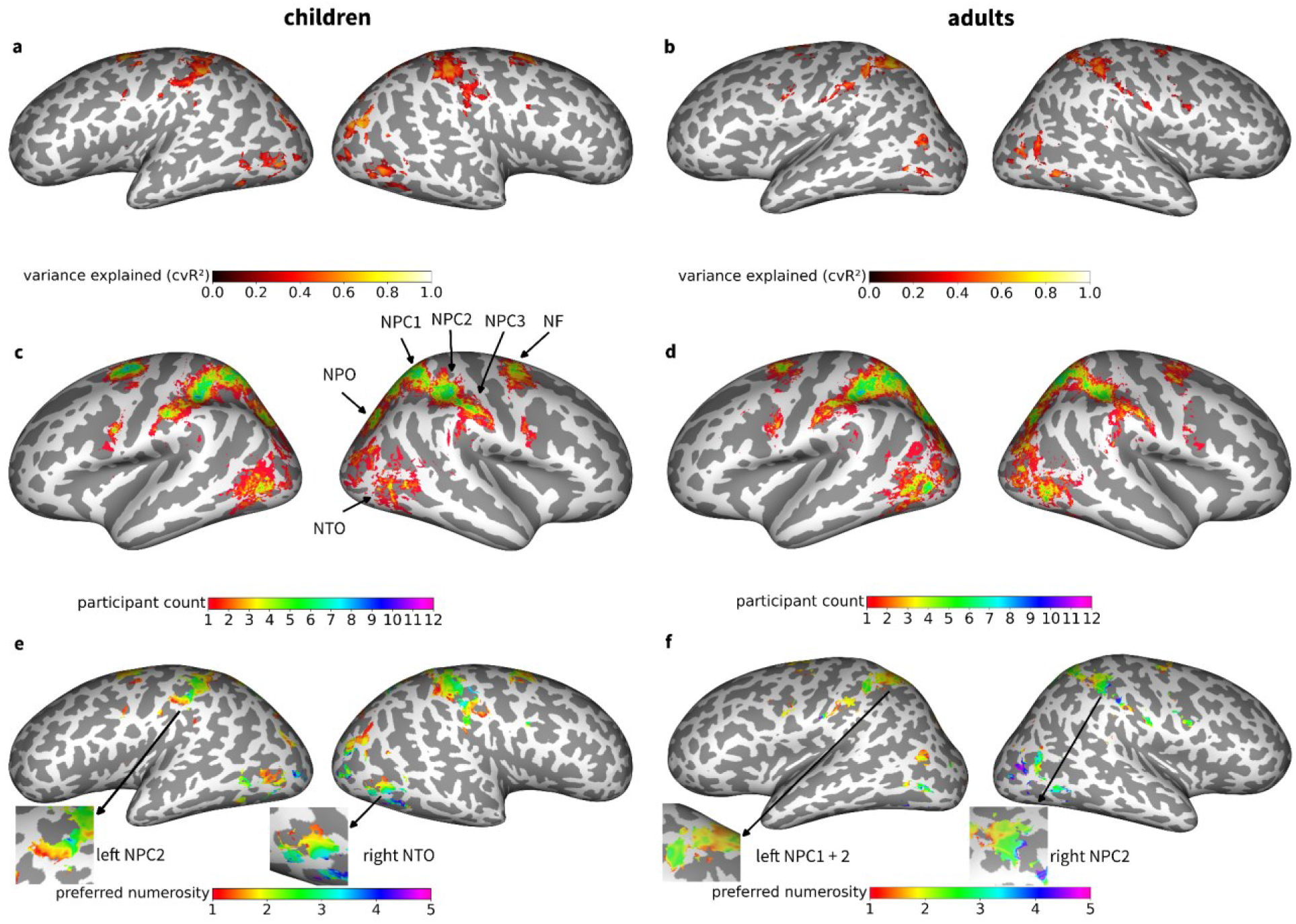
Spatial distribution of hemodynamic responses to visual numerosity. Inflated surface maps showing regions in which neural tuning models explain a significant amount of the variance for **a** an example child and **b** an example adult subject (same subjects as in Fig. 2). The colorbar indicates the variance explained by the model. **c,d**, Participant count maps for **c** children and **d** adults in standard space (FreeSurfer fsaverage). The colorbar indicates how many subjects exhibit tuning to numerosity according to the variance explained criterion (p < 0.05, Bonferroni-corrected). **e,f**, Preferred numerosity maps of **e** the child and **f** the adult (same subjects as in a,b). The colorbar indicates the estimated preferred numerosity. Inset maps show exemplary topographic organization in one numerosity map per hemisphere and subject. NTO = temporo-occipital visual numerosity field; NPO = parieto-occipital visual numerosity field; NPC1,2,3 = parietal visual numerosity fields; NF = frontal visual numerosity field.

**Fig. 4.**
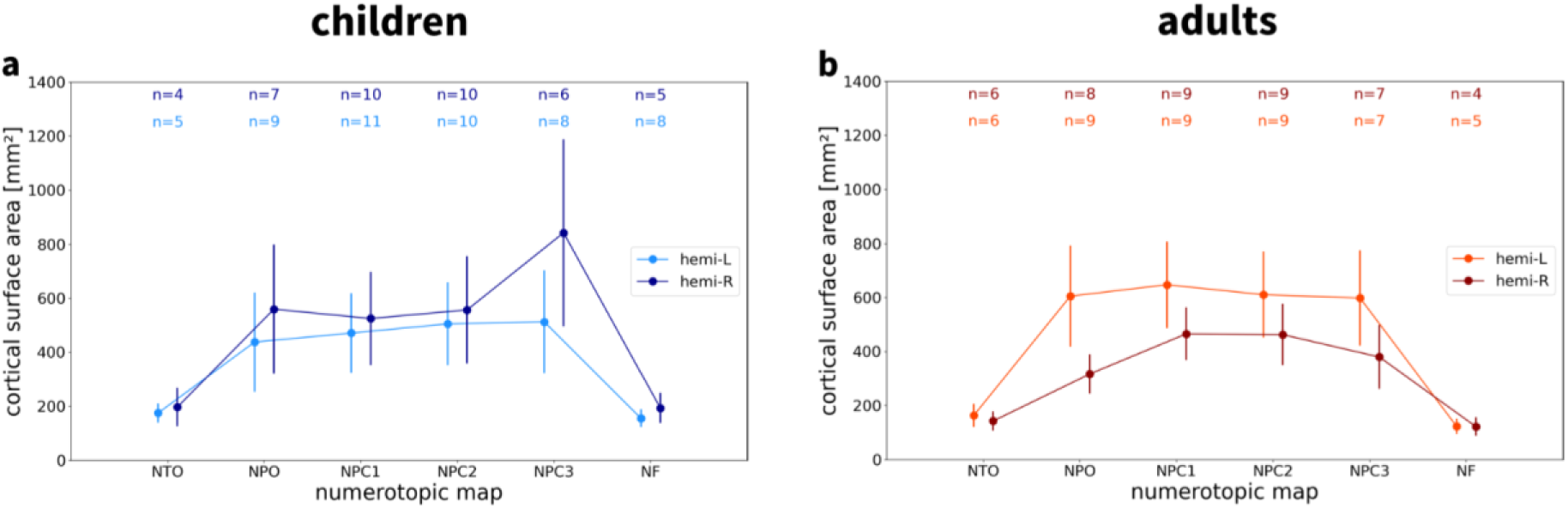
Surface area of numerosity maps. Cortical surface areas of **a** children and **b** adults are shown separately for each hemisphere and topographic map. Numbers on top of the graph (n) indicate the number of subjects which exhibit each map, separated by hemisphere, based on minimum cluster size and maximum distance from the center of the map across the group (see Methods). NTO = temporo-occipital visual numerosity field; NPO = parieto-occipital visual numerosity field; NPC1,2,3 = parietal visual numerosity fields; NF = frontal visual numerosity field L = left hemisphere; R = right hemisphere.

### Topographic organization of visual numerosity maps

For each single subject, supra-threshold clusters were assigned to topographic maps if their cortical surface area was larger than a minimum area (*A*_*min*_ = *50 mm^2^*) and if their distance from previously reported center coordinates^12^ was smaller than a maximum distance (*d*_*max*_ = *25 mm*). Analyzing the cortical surface extent of those maps, we found that most topographic maps were identified in the majority of children and adults (Fig. 4).

For each hemisphere, we then partitioned the preferred numerosity of all vertices into bins with a width of 0.5. This resulted in eight bins (1-1.5, 1.5-2, 2-2.5, 2.5-3, 3-3.5, 3.5-4, 4-4.5, 4.5-5) and corresponding bin centers (1.25, 1.75, 2.25, 2.75, 3.25, 3.75, 4.25, 4.75). Next, we calculated how much cortical surface is devoted to representing each level of preferred numerosity. We found a negative correlation between cortical surface area and preferred numerosity (Fig. 5; for replication in other subjects, see Supplementary Fig. S5), indicating that fewer cortical surface area represents higher numerosities. Simulations confirmed that, for numerosity-selective voxels, preferred numerosity is accurately estimated by the tuning model, as long as it falls into the presented stimulus range. These results exclude a model bias towards smaller preferred numerosities which could be assumed given that low-numerosity vertices (with a preferred numerosity smaller than around 2.5) were more frequently observed than medium-numerosity vertices (between 2.5 and 5).

**Fig. 5.**
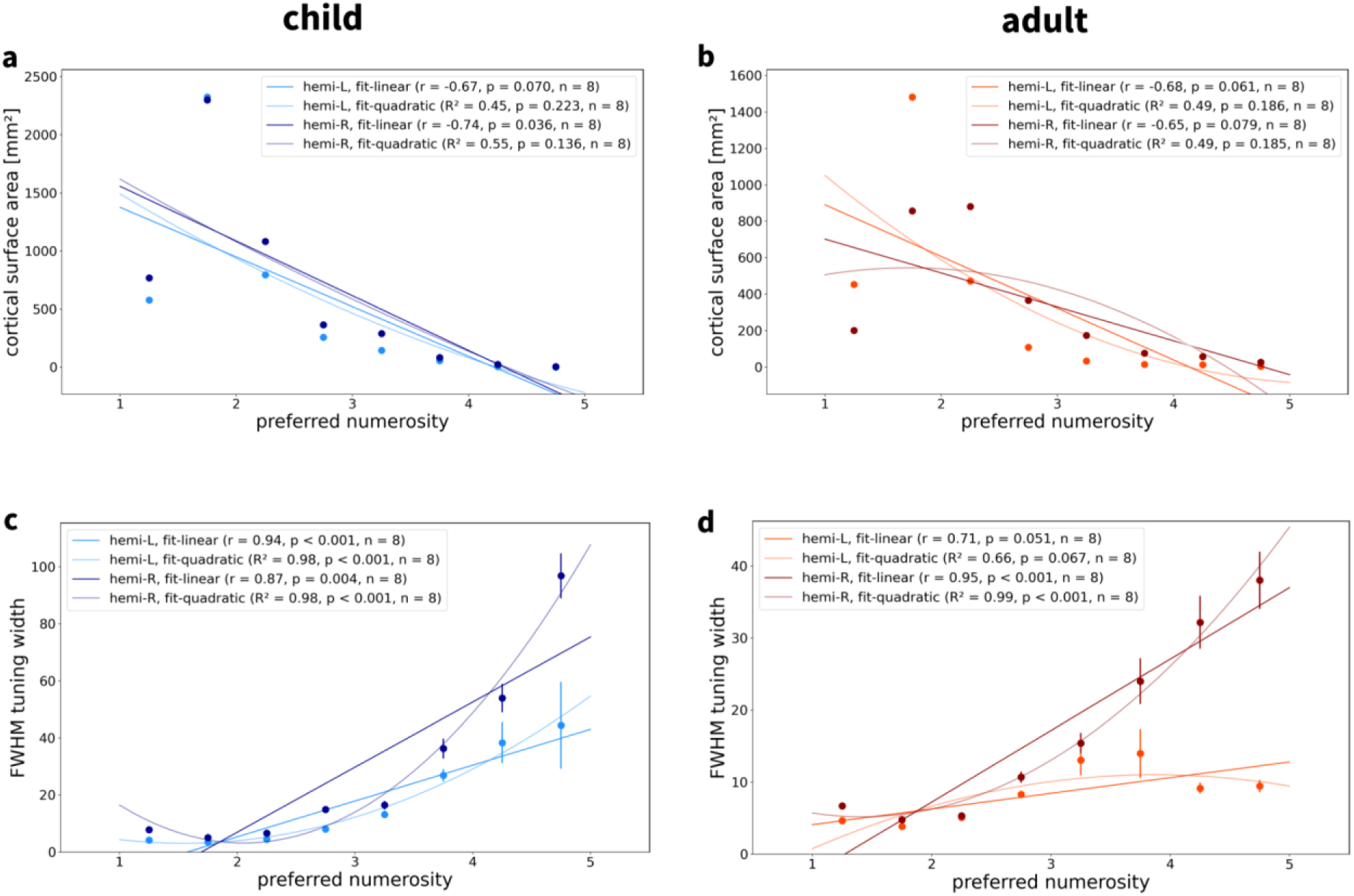
Associations between preferred numerosity, surface area and tuning width. **a,b**, Associations between preferred numerosity and cortical surface area in all supra-threshold vertices for **a** an example child and **b** an example adult (same subjects as in Fig. 2 and Fig. 3). Points correspond to the summed cortical area of all surface triangles of vertices with an average preferred numerosity falling in the respective range (bin width = 0.5). **c,d**, Relationship between preferred numerosity and FWHM tuning width in all supra-threshold vertices for **c** visual numerosity and **d** auditory numerosity (same subjects as in **a,b**). Points correspond to mean FWHM in each preferred numerosity bin. Error bars correspond to the standard error of the mean. Solid lines correspond to simple linear regression fits or fitted quadratic curves. L = left hemisphere; R = right hemisphere; r = Pearson correlation coefficient; R² = variance explained; n = number of numerosity bins.

We also extracted vertex-wise tuning width, measured as full width at half maximum (FWHM), and analyzed its relationship with binned preferred numerosity in each hemisphere. This analysis revealed a positive correlation between preferred numerosity and FWHM tuning width (Fig. 5; for replication in other subjects, see Supplementary Fig. S6). Accordingly, the larger the numerosity to which a population responds most strongly, the higher the extent to which it also responds to neighboring numerosities.

Next, we ran linear mixed-effects models, using summed cortical surface area or average FWHM tuning width as the dependent variable, binned preferred numerosity and brain hemisphere as independent variables while modeling subject as a random effect. We detected significant negative effects of preferred numerosity on surface area (all p < 0.001) and significant positive effects of preferred numerosity on tuning width (all p < 0.001) in both hemispheres (Table 1), indicating that the larger the numerosity, the smaller the cortical surface area covered and the wider the tuning. There were significant effects of hemisphere on surface area in adults (p = 0.065), but not in children (p = 0.775), and no significant effects of hemisphere on tuning width (children: p = 0.738; adults: p = 0.497). In adults, more cortical surface was devoted to numerosity in the left hemisphere (Table 1; also see Fig. 4).

**Table 1.**
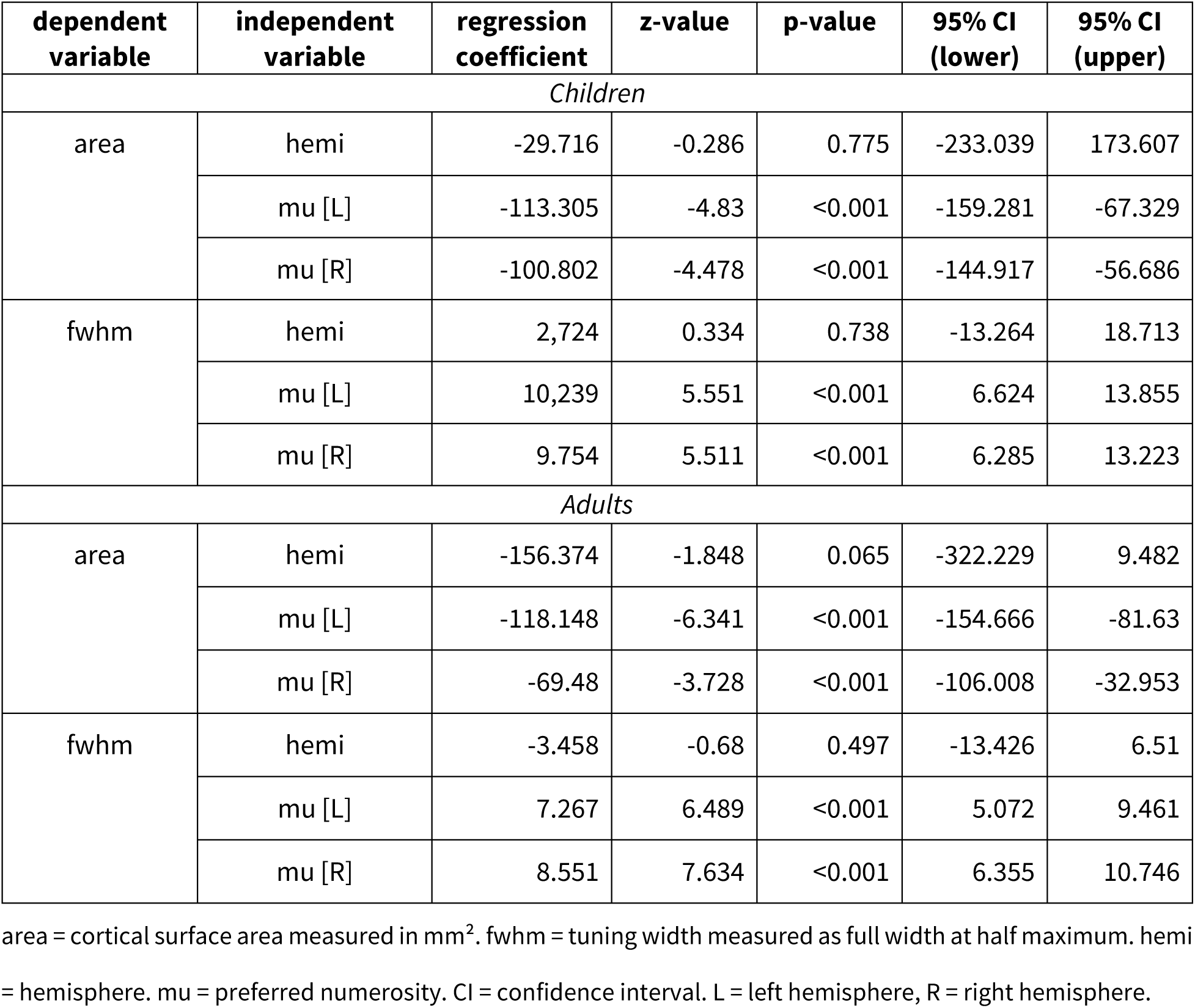
Associations of surface area and tuning width with preferred numerosity.

| dependent variable | independent variable | regression coefficient | z-value | p-value | 95% CI (lower) | 95% CI (upper) |
| --- | --- | --- | --- | --- | --- | --- |
| *Children* |  |  |  |  |  |  |
| area | hemi | -29.716 | -0.286 | 0.775 | -233.039 | 173.607 |
|  | mu [L] | -113.305 | -4.83 | <0.001 | -159.281 | -67.329 |
|  | mu [R] | -100.802 | -4.478 | <0.001 | -144.917 | -56.686 |
| fwhm | hemi | 2,724 | 0.334 | 0.738 | -13.264 | 18.713 |
|  | mu [L] | 10,239 | 5.551 | <0.001 | 6.624 | 13.855 |
|  | mu [R] | 9.754 | 5.511 | <0.001 | 6.285 | 13.223 |
| *Adults* |  |  |  |  |  |  |
| area | hemi | -156.374 | -1.848 | 0.065 | -322.229 | 9.482 |
|  | mu [L] | -118.148 | -6.341 | <0.001 | -154.666 | -81.63 |
|  | mu [R] | -69.48 | -3.728 | <0.001 | -106.008 | -32.953 |
| fwhm | hemi | -3.458 | -0.68 | 0.497 | -13.426 | 6.51 |
|  | mu [L] | 7.267 | 6.489 | <0.001 | 5.072 | 9.461 |
|  | mu [R] | 8.551 | 7.634 | <0.001 | 6.355 | 10.746 |

area = cortical surface area measured in $\text{mm}^2$ . fwhm = tuning width measured as full width at half maximum. hemi = hemisphere. mu = preferred numerosity. CI = confidence interval. L = left hemisphere, R = right hemisphere.

In the subsequent step, we fitted models describing observed preferred numerosities based on each vertex’s pial surface coordinates in standard space. These models used polynomial expansion of x-, y- and z-coordinates up to order 5 and were estimated with 10-fold cross-validation (see Methods). This approach resulted in predicted preferred numerosities which were compared to actual preferred numerosities across vertices. For all maps, there were significant positive correlations between predicted and actual numerosities (Fig. 6a,b; for replication in other subjects, see Supplementary Fig. S7). Hence there is a relationship between cortical surface coordinates and represented numerosity which, however, is not necessarily linear. Finally, we analyzed the range of numerosities within each topographic map on the cortical surface in both hemispheres and found that not all maps cover the whole range of numerosities presented during the experiment (Fig. 6c,d; for replication in other subjects, see Supplementary Fig. S8).

**Fig. 6.**
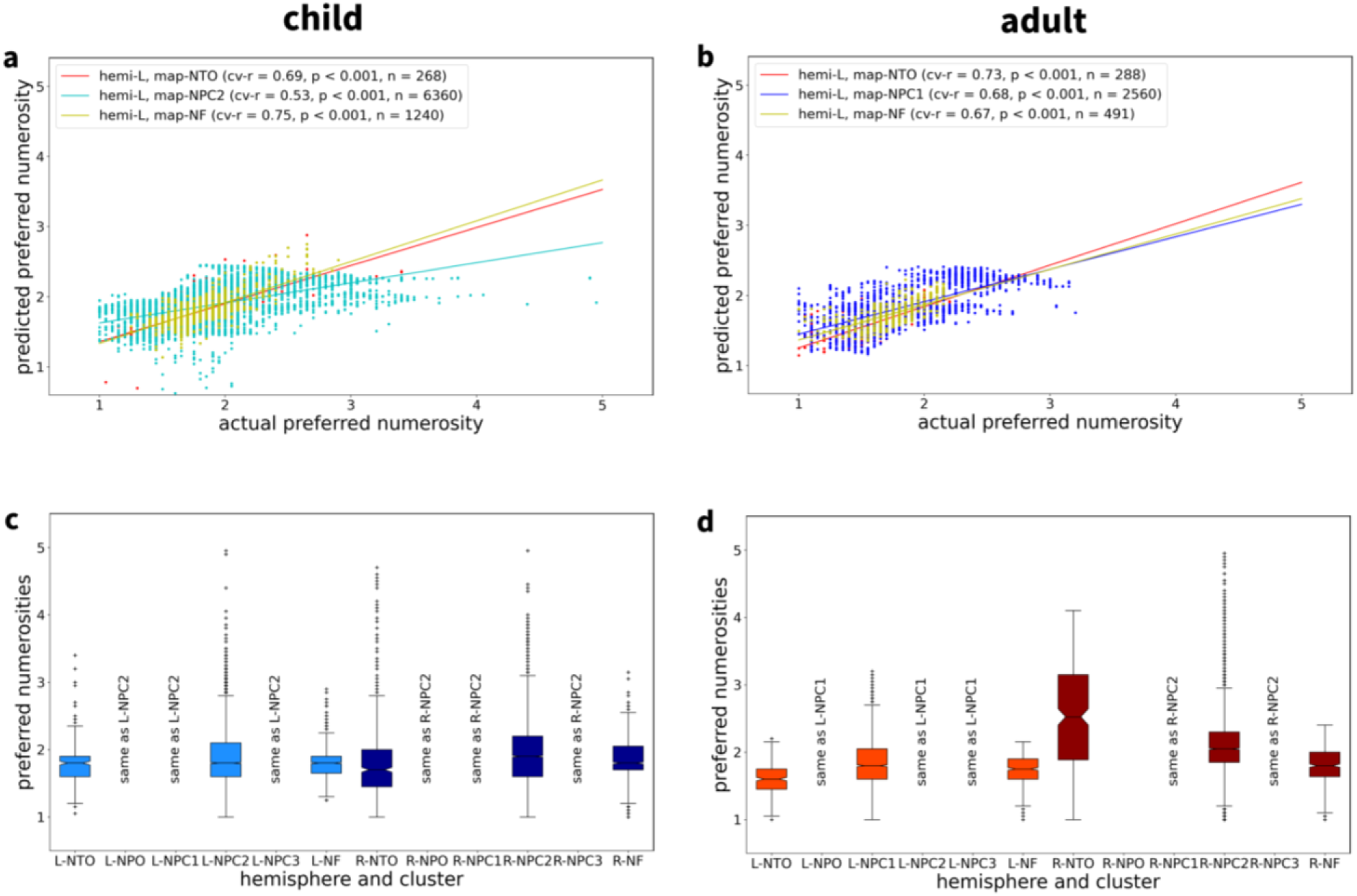
Progression of preferred numerosity within topographic maps. **a,b**, Plots of preferred numerosity, predicted from cortical surface coordinates, against actual numerosity for left-hemisphere maps of **a** one child and **b** one adult example subject (same single subjects as in Fig. 2, 3 and 5). Significant predictive correlations (cv-r) indicate that preferred numerosity can be predicted from surface coordinates, suggesting a spatially systematic organization of responses to numerosity. **c,d**, Ranges of represented numerosities for all topographic maps found in each example subject. Whenever a cluster covered multiple maps in one hemisphere (e.g. NPC1,2,3), it was assigned to the map with the closest center (described with same-as labels). L = left hemisphere; R = right hemisphere; NTO = temporo-occipital visual numerosity field; NPO = parieto-occipital visual numerosity field; NPC1,2,3 = parietal visual numerosity fields; NF = frontal visual numerosity field; cv-r = predictive correlation; n = number of vertices in a map.

## Discussion

We found a receptive field-like neural population code underlying visual quantity detection in individual developing human brains. Population coding for numerosity was revealed by electrophysiologically inspired models applied to hemodynamic responses recorded with 7 Tesla high-field magnetic resonance imaging. We identified six bilateral topographic maps with similar Gaussian tuning properties in most children and adults. At the same time, there were pronounced individual differences regarding the location of the maps and the variance explained. These effects were unrelated to attentional task performance.

The observation that receptive field models explained comparable amounts of hemodynamic variance in children and adults is consistent with the interpretation that population coding for small numerosities has already reached a similar level of precision in late childhood compared to adulthood. This finding is remarkable in the context of behavioral studies suggesting that number discrimination ability still refines substantially from adolescence to adulthood^4^ and that, even in adolescence, children have less sustained attention capacities than adults^14^. It is thus plausible to assume that developmental cognitive refinement in late childhood is unrelated to neural coding for small numerosities. This prediction remains to be tested empirically by means of a precision imaging approach following children longitudinally towards adulthood^15^.

An important open question is how tuned topographic numerosity maps emerge in the course of early brain development. It is possible that at least some of these maps already form in the first months of life, given the behavioral evidence that numerosity discrimination ability increases from a 1:3 ratio at birth to a 2:3 ratio at 10 months of age^3^. Due to major feasibility concerns, however, we decided against putting this assumption to the test. One critical challenge is that high-field fMRI at a field strength of 7 Tesla is particularly susceptible to strong head motion which is practically unavoidable in unsedated infants, toddlers and young children^16,17^. Even if head motion could be prevented, the amount of data that could be collected in a single session would on average be as low as about 5 minutes in infants and about 10 minutes in toddlers^16^. Accordingly, to generate a functional dataset comparable to the dataset collected in the present study (i.e., 41 minutes) infants would have to undergo at least eight tightly scheduled scanning sessions in an awake state which seems virtually impossible. Crucially, moving from 7 Tesla to 3 Tesla would not be a viable alternative since it is known that one run at 3 Tesla has, on average, only about a quarter of the model predictive power of one run at 7 Tesla^18^. This means that the numerosity mapping experiment conducted in the present study would require almost 3 hours of high quality 3 Tesla data which is infeasible in infants and toddlers and also unrealistic for young children.

Numerosity co-varies with non-numerical physical features of a stimulus. Specifically, varying the number of items in a set changes spatial features while fixing one feature causes another to vary with the number of items^13^. Responses to numerosity should thus be comparable across systematically varied spatial features to be considered as abstract^19^. In line with this criterion for abstract representation, responses to visual numerosity have been consistently shown to be equal across co-varying spatial features, such as total area, size, density, circumference and shape^13^. The robustness of this effect has been replicated across electrophysiological and hemodynamic recording techniques both in human and nonhuman primates. Accordingly, to keep the experimental effort within a reasonable limit for children we did not create additional conditions in which other features than total area are kept constant since each feature would have required running an additional experiment with a duration of 40 minutes. Whether the current results generalize to other co-varying spatial features beyond total area remains to be evaluated in future investigations.

## Methods

### Participants

Following previous electrophysiological and magnetic resonance imaging work, we employed a single-subject design focused on individual human datasets^10-13^. Fifteen children (aged 11–12 years, all right-handed, eight males) and 12 adults (aged 20–34 years, all right-handed, six males) were recruited to demonstrate reproducibility and generalization across subjects and age groups (see Supplementary Table S1). Three children were excluded from further data analysis since less than 75% of their runs passed our quality control procedure (see functional preprocessing section below). Given that the datasets were in agreement and led to similar conclusions with high statistical confidence, we did not consider it as scientifically necessary, economically reasonable and ethically plausible to include more participants. All participants had normal or corrected to normal visual acuity. Children were in the fifth grade of school. Adults were well educated (eleven college degrees, one vocational degree). All participants gave written informed consent to participate. The study was approved under the reference 317/19-ek by the Ethics Committee at the Medical Faculty of the University of Leipzig, Germany (IRB00001750).

### Sensory stimulation

#### Visual stimuli

In the visual experiment, stimuli were projected on a 34 × 25 cm screen inside the MRI bore with a resolution of 1,024 × 768 pixels. Participants viewed this screen through a mirror attached to the head coil. Stimuli were black dots on a middle gray background presented at the center of the display. To guide fixation and maintain attention, dots appeared within a radius of 0.75° (visual angle) of a diagonal cross of thin red lines covering the entire display throughout the experiment. To control for low-level non-numerical visual features, summed surface area (and thus luminance) was kept constant and dots were distributed homogeneously (preventing perceptual grouping) while randomly varying the positions of the dots. All stimuli were created using PsychoPy version 2021.2.3^20^. Other low-level features have been repeatedly and consistently shown not to influence numerosity-specific hemodynamic time courses^12,13^.

#### Stimulation procedure

Numerosities one through five were used to capture both the subitizing range (one through three) and the estimation range (four to five)^21,22^. They were first presented in ascending order in blocks changing every 4,200 ms. Within each block of the visual experiment, a dot array was shown six times for 300 ms each, alternating with 400 ms of gray background. Short stimulus durations were chosen to prevent participants from counting and to avoid adaptation effects. Short equal pauses were introduced to prevent participants from keeping numerosities in working memory. After the ascending numerosity sequence, twenty items were presented 24 times for 16.8 s including pauses. Next, numerosities one through five were presented again, but in descending order, followed by another block of twenty items. This stimulation cycle (1 – 2 – 3 – 4 – 5 – 20 – 5 – 4 – 3 – 2 – 1 – 20) was repeated four times in each run. The rationale for the long baseline period was that hemodynamic responses could return to baseline for numerosity-selective regions with small preferred numerosities and little neural responses to 20 items. The rationale for the ascending-descending numerosity presentation was to avoid adaptation effects, because each stimulus is preceded by both lower and higher numerosities in different blocks of presentation, thereby counterbalancing potential effects of previously presented numerosity^13^. Additionally, keeping this fixed and smooth order of presentation is known to result in more homogeneous topographic maps than a pseudorandom stimulation sequence. Alternating between ascending and descending presentation served to minimize neural adaptation by ensuring each stimulus was preceded by a stimulus inducing both a lower and higher response. Stimuli were presented with PsychoPy version 2021.2.3^20^. There were eight runs for a total experiment duration of 41 min.

#### Target detection task

In 10% of all stimulus events, white instead of black dots were shown. Participants were instructed to respond to these events by pressing a button. This simple target detection task was introduced to ensure participants were paying attention and following the task.

### Data acquisition

#### Magnetic resonance imaging

T2*-weighted functional magnetic resonance imaging data were acquired with a 1-channel transmit / 32-channel receive coil (NOVA Medical, Wilmington MA, USA) on a 7 Tesla Magnetom TERRA scanner (Siemens Healthineers, Erlangen, Germany). The scanner was operated in the CE-certified clinical mode. We used a 2D gradient-echo echo-planar imaging (2D-EPI) sequence for acquiring 41 slices (thickness = 1.75 mm) with a field of view FOV = 192 mm x 192 mm and a matrix size = 110 x 110, resulting in an isotropic voxel size of 1.75 mm x 1.75 mm x 1.75 mm. The slice package had a transversal orientation, but was tilted towards coronal orientation in order to spare anterior frontal and temporal lobes where pronounced B0 and B1 inhomogeneities degrade image quality at 7T. To achieve a repetition time TR of 2.1 s, the acquisition was accelerated using GRAPPA with an iPAT factor of 3. An echo time TE of 24 ms, a flip angle FA of 70° and a receiver bandwidth of 1684 Hz/Px was used. 145 volumes were acquired per run resulting in a single run time of 05:04,50 min. For distortion correction, an EPI data set using the same parameters, but with the phase-encoding direction flipped from anterior-posterior to posterior-anterior was acquired. For anatomical referencing, a T1-weighted image was acquired using an MP2RAGE sequence^23^ with the following parameters: TR = 5.000 ms, TE = 2.01 ms, TI_1/2_ = 900 ms / 2.750 ms, FA_1/2_ = 5° / 3°, isotropic voxel size of 0.7 mm x 0.7 mm x 0.7 mm.

### Data preprocessing

Preprocessing was performed using FreeSurfer 7.4.1^24^ and fMRIprep 23.1.4^25^, based on Nipype 1.8.6^26^. The following description of fMRIPrep’s preprocessing is based on a boilerplate distributed with the software covered by a ‘no rights reserved’ (CC0) license. For more details about the pipeline, see the section corresponding to each workflow in the fMRIPrep documentation.

#### Structural preprocessing

Before running fMRIPrep, we segmented the MP2RAGE UNI images using a custom workflow implemented in the python package fmritools 1.0.3 (https://doi.org/10.5281/zenodo.10573042). MP2RAGE image backgrounds were denoised as described in O’Brien et al. 2014^27^ and the resulting images were bias-corrected with SPM12 (https://www.fil.ion.ucl.ac.uk/spm/software/spm12). Then the first five Freesurfer cortical reconstruction steps were applied before computing a skull-strip mask based on the INV2 of the MP2RAGE image. This skull-strip mask was applied to the remaining Freesurfer cortical reconstruction pipeline. As a first step in fMRIPrep, the denoised and thresholded structural images were corrected for intensity non-uniformity (INU) with N4BiasFieldCorrection^28^, distributed with ANTs^29^. Images were then skull-stripped with a Nipype implementation of the antsBrainExtraction.sh workflow (implemented in ANTs), using OASIS30ANTs as target template. Brain tissue segmentation of cerebrospinal fluid (CSF), white-matter (WM) and gray-matter (GM) was performed on the brain-extracted images using FSL FAST^30^. In the case that two T1w images were available for a subject, a structural T1w reference map was computed after co-registration of the INU-corrected T1w images using mri_robust_template implemented in FreeSurfer 7.3.2^31^. Otherwise, the INU-corrected T1w image was used as T1w reference throughout the pipeline. Brain surfaces were reconstructed using the recon-all procedure of FreeSurfer 7.3.2^24^. Brain masks estimated before were refined based on Mindboggle’s method to integrate ANTs-derived and FreeSurfer-derived segmentations of the cortical gray matter. This way, the best results of the ANTs-derived and FreeSurfer-derived cortical gray-matter segmentations could be included in the final masks^32^.

#### Functional preprocessing

Before functional data preprocessing, image quality was assessed using MRIQC and high-motion runs (framewise displacement >1.7mm for >10% of volumes) were excluded from further analysis^33^. Using fMRIPrep 23.1.4, for each of the up to 8 functional runs per subject, the following preprocessing steps were taken: First, a reference volume for estimating head-motion parameters and its skull-stripped version were generated. Head motion parameters (transformation matrices, and six corresponding rotation and translation parameters) were estimated before any spatiotemporal filtering using FSL MCFlirt^34^. B0-nonuniformity maps (fieldmaps) were generated from two (or more) EPI references with FSL topup^35^. Fieldmaps were then aligned with rigid registration to the target EPI reference run. Field coefficients were mapped onto the reference EPI using the transformation matrices generated when aligning the fieldmaps with the reference EPI. Next, functional runs were slice-time-corrected to 1.025 s (which is equivalent to 0.5 of the slice acquisition range of 0-2.05 s) using AFNI’s 3dTshift^36^.

Functional reference images were then co-registered to the T1w reference images by applying boundary-based registration with bbregister in FreeSurfer^37^. Co-registration was configured with six degrees of freedom. Several confounding time-series were calculated based on the preprocessed functional images: framewise displacement (FD), DVARS (D referring to temporal derivative of time courses, VARS referring to root mean square variance over voxels) and three region-wise global mean signals extracted from within the CSF, the WM, and the whole-brain masks. FD was computed using two formulations based on the absolute sum of relative motions and the relative root mean square displacement between affines^34,37^. FD and DVARS were calculated for each functional run using their implementations in Nipype which follow the definitions by Power and colleagues^38^.

These nuisance variables and head motion estimates were returned as a confounds file for each individual run. Specifically, the confound time series derived from head motion estimates and global signals were expanded by including temporal derivatives and quadratic terms for each^39^. Frames that exceeded a threshold of 1.7 mm FD or 1.5 standardized DVARS were annotated as motion outliers. Additional nuisance time series were calculated by performing principal components analysis for the signals found within a thin band (crown) of voxels around the edge of the brain, as proposed by Patriat and colleagues^40^.

The functional time-series were resampled onto the fsnative (individual participant) and fsaverage (standard space) surfaces following the FreeSurfer reconstruction nomenclature. All resamplings were performed with a single interpolation step by composing all the pertinent transformations (i.e. head-motion transform matrices, susceptibility distortion correction if available, and co-registrations to anatomical and FreeSurfer output spaces). Non-gridded (surface) resamplings were performed using mri_vol2surf in FreeSurfer. No spatial or temporal smoothing was applied^41^.

### Numerosity analysis

#### Numerosity population receptive field models

Task-based fMRI data were analyzed using a well-established forward modelling approach^12,13,42^. Each vertex is thought to represent the presented numerosities using a logarithmic Gaussian tuning curve characterized by two parameters: the preferred numerosity *μ* (the numerosity to which the population responds strongest) and a tuning width *ω* (the full width at half maximum (FWHM) of the response function). Given presented numerosity *x*_*t*_ at continuous time *t*, the neuronal response at *t* is given by

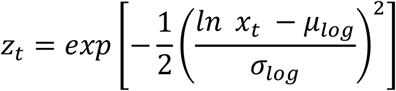

where *μ*_*log*_ =*ln μ* and *σ*_*log*_ is the standard deviation of the response function in logarithmic numerosity space from which the FWHM tuning width in linear numerosity space can be calculated:

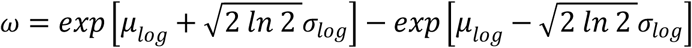

Given an assumed hemodynamic response function (HRF) ℎ(*t*), the hemodynamic response is given by the convolution of the neuronal response with the HRF:

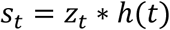

Finally, a linear relationship is assumed between this hypothesized hemodynamic response, governed by the neural tuning parameters *μ* and *ω*, and the preprocessed fMRI signal

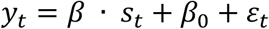

where *β* is a scaling factor of the hypothesized signal, *β_0_* is the baseline level of hemodynamic activity and *ɛ*_*t*_ are noise terms independent and identically distributed (i.i.d.) with zero mean and unknown variance. Before estimation of tuning parameters *μ*, *ω* and *β*, preprocessed fMRI signals were standardized to units of percent signal change and 12 confound variables were regressed from standardized signals (six motion parameters; WM, CSF and global signal; three cosine regressors modelling temporal drifts). Then, signals were averaged across runs (but not cycles) and numerosity analysis was performed for a single averaged run.

#### Estimation of numerosity tuning parameters

The numerosity pRF model defines a likelihood function of the measured data *y*, given the tuning parameters *μ* and *ω*. Some combinations of tuning parameters make the measured signals more likely than others and the goal is to find the tuning parameters which are most compatible with the measured fMRI data. To this end, we specified a large grid of plausible values: *μ* from 0.8 to 5.2 in steps of 0.05, *σ*_*log*_ from 0.05 to 30 in steps of 0.05 (such that *0*.*12* ≤ *ω* ≤ *34*.*17* for the lowest presented numerosity *μ* = *1* and *0*.*59* ≤ *ω* ≤ *170*.*9* for the highest presented numerosity *μ* = *5*; cf. equation above). For each possible combination *μ* and *ω*, the ensuing neuronal responses *z* were calculated using the sequence of presented numerosities *x* and the ensuing hemodynamic signals *s* were generated using the canonical HRF, as implemented in SPM. Then, *β* was estimated from the measured fMRI signal *y* using ordinary least squares, resulting in a single log-likelihood value for each combination of *μ* and *ω*:

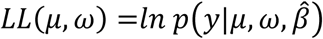

Finally, the optimal tuning parameters are those which maximize the log-likelihood function across all possible combinations:

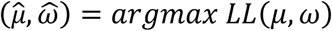

When not accounting for serial correlation in fMRI signals, this is equivalent to estimating tuning parameters by minimizing the residual sum of squares (RSS), as was done previously^4,7^.

#### Cross-validated estimation of model fit

For unbiased estimation of tuning model precision, we applied split-half cross-validation and partitioned recorded fMRI data into odd runs (runs 1, 3, 5, 7) and even runs (runs 2, 4, 6, 8) for each participant. Then, tuning parameters *μ* and *ω* were estimated from one half of the data (averaged over runs) and used to generate expected hemodynamic time courses. These time courses were then compared against the measured signals from the other half of the data (averaged over runs), yielding a coefficient of determination (*R^2^*). Cross-validated model fit was obtained by averaging coefficients of determination from odd and even runs. This is referred to as cvR² and was used as the primary criterion for selecting supra-threshold vertices.

#### Exploration of alternative analysis options

Parameter estimation via maximum likelihood is equivalent to ordinary least squares under independent and identically distributed errors and no serial correlations. We did not account for serial correlations using e.g. restricted maximum likelihood and weighted least squares, since this did not improve tuning parameter estimates in a pre-analysis simulation study, even if serial correlations were present in the simulated signals. We also did not include hemodynamic derivatives to account for possible variation of HRF between subjects and vertices, because this was found to generate rather noisy numerosity-selective clusters on the cortical surface in a single-subject pilot analysis.

Finally, we also tested linear tuning functions in which the neuronal response at *t* is given by

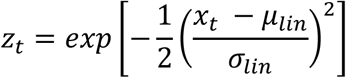

where *μ*_*lin*_ = *μ* and *σ*_*lin*_ is the standard deviation of the response function in linear numerosity space from which the FWHM tuning width can be calculated:

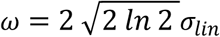

Example results obtained from analyses based on linear tuning functions are provided in Supplementary Figure S9.

#### Testing for categorical numerosity effects

To rule out the possibility that neuronal populations merely react to the categorical difference between small numerosities (1-5) and the large numerosity (20), we implemented a vertex-wise classical GLM analysis. At the single-subject level, a categorical model with six HRF regressors for numerosities 1, 2, 3, 4, 5, 20 and twelve confound variables (see above) was specified and estimated. For each subject, a contrast map “1-5 minus 20” was calculated using the contrast vector 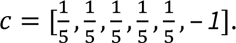 At the group level, a one-sample t-test was run across contrast maps from all subjects and statistical inference was performed using t-contrasts for positive effects (1-5 > 20) and negative effects (20 > 1-5) of small numerosities.

Except for some parietal clusters, these effects were restricted to higher responses for the large numerosity (20 > 1-5) (see Supplementary Figure S10). However, observing such effects when using the numerosity tuning model would require either a preferred numerosity close to this stimulus level (*μ* ≈ *20*) or a small preferred numerosity along with negative scaling factor (*β* < *0*). Since both are filtered out when selecting numerosity-selective vertices (requiring positive scaling factor and numerosity between 1 and 5; see below), our results are unlikely to reflect categorical differences between small and large numerosities. Moreover, theoretical simulation analyses indicate that differential responses to numerosities can emerge even with high tuning width (e.g. fwhm ≈ 80) for both, low and medium preferred numerosities (see Supplementary Figure S11).

### Statistical analyses

#### Extraction of parameter estimates

The numerosity pRF model returns four quantities for each vertex: estimated preferred numerosity 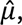 estimated FWHM tuning width 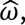 estimated scaling factor 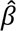 and cross-validated *R^2^* (cvR²). Only vertices that exhibited a positive scaling factor 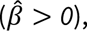 an estimated preferred numerosity inside the presented stimulus range 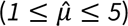 and statistically significant variance explained (p < 0.05, Bonferroni-corrected; see below) were included in the following analyses. The rationale for excluding vertices with negative scaling factor was that the preferred numerosity could not be interpreted as the stimulus generating the maximum response. The rationale for excluding vertices with preferred numerosity outside the stimulus range is that their tuning functions will be monotonically increasing or decreasing throughout the stimulus range and thus not display actual tuning. Simulations indicate that numerosity-selective voxels can be distinguished from no-signal voxels with high sensitivity (≈ 85%) and specificity (≈ 100%) and that preferred numerosity is accurately estimated, if it falls into the presented stimulus range (https://github.com/SkeideLab/EMPRISE-KIDS/blob/main/code/Python/Demo.pdf).

#### Inferential statistics for variance explained

To assess statistical significance of variance explanation by the tuning model, we performed an F-test of the model including the numerosity regressor (generated using tuning parameters obtained from the other half of the data) against a null model including only the baseline regressor:

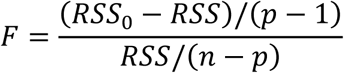

where *n* = *145* is the number of scans per run and *p* = *2* is the number of free parameters in the numerosity pRF model (*β_0_*, *β*). Note that *μ* and *ω* were not counted into the number of free parameters, because they were estimated from independent data (the respective other half of the fMRI runs). Under the null hypothesis *H_0_*: *R^2^* = *0*, this test statistic is following an F-distribution with *p* − *1* numerator and *n* − *p* denominator degrees of freedom. The F-statistic can also be expressed in terms of (cross-validated) variance explained:

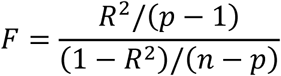

Based on this statistic, cvR² maps were thresholded by applying a significance level of *α* = *0*.*05*, Bonferroni-corrected for the number of vertices in the current hemisphere. For example, with 100,000 vertices per hemisphere (actual numbers ranged between 84,725 and 110,511 for children as well as 90,915 and 116,566 for adults in native subject space), this implies a variance explained threshold of *R^2^* = *0*.*1625*. Only vertices surpassing this threshold (derived from *n*, *p*, *α* and number of vertices) were retained for the following analyses.

#### Calculation of cortical surface area

Alongside with estimated tuning parameters, the coordinates on the pial surface in native subject space were extracted for each supra-threshold vertex. We here chose pial surface rather than white matter or midthickness surface coordinates, as these most precisely correspond to actual cortical surface extents in native subject space. After filtering numerosity maps with the criteria mentioned above, AFNI’s SurfClust function was used to form clusters of vertices with an edge distance of at most 1 on the cortical surface (i.e. vertices needed to be directly connected to belong to the same cluster). For each cluster, the total area on the cortical surface was calculated as the summed area of all triangles for which all three nodes belonged to the cluster. Only clusters having a minimum cortical surface area *A*_*min*_ = *50 mm^2^* were retained.

#### Identification of topographic maps

In order to investigate topographic organization in larger numerosity-selective clusters, supra-threshold clusters were assigned one of the previously reported (Harvey & Dumoulin, 2017) six topographic maps (temporo-occipital, NTO; parieto-occipital, NPO; in parietal cortex, NPC1/2/3; in prefrontal cortex, NF; N = numerosity) by being in less than some distance (*d*_*max*_ = *25 mm*) from previously reported center coordinates^12^ with at least one vertex in standard space.

#### Visualization of numerosity selectivity

The estimated tuning parameters 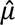 and 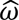 describe a logarithmic Gaussian tuning function and an expected time course of hemodynamic activity in a single cycle of numerosity presentation. These are plotted for a low-numerosity vertex 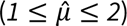 and a high-numerosity vertex 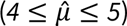 from exemplary single subjects (see Figure 2 and Supplementary Figures S1/S2). In addition to this, vertex-wise variance explained and vertex-wise preferred numerosity are visualized in native subject space for all vertices passing the above filtering criteria (see Figures 3a/b and 3e/f and Supplementary Figures S3/S4). For each vertex in standard space, the number of subjects showing numerosity selectivity according to these criteria are shown as participant count maps (see Figure 3c/d).

#### Statistical analysis of topographic organization

For each of the six topographic maps in each hemisphere, we calculated the average cortical surface area (see Figure 4) as well as the represented range of preferred numerosities (see Figure 6c,d and Supplementary Figure S8). Additionally, we investigated whether there is a topographic organization within those maps, i.e. progression of preferred numerosity along the cortical surface. Previously, this was done by manually delineating start, end and borders of topographic maps, establishing distance along the so-defined principal axis of a topographic map and calculating average preferred numerosity as a function of binned cortical distance^12,13^. As manual delineation might lead to non-reproducible results and would be tedious given the amount of data (24 participants, six maps each), we instead used a linear regression model with preferred numerosity as the dependent variable and pial x-, y- and z-coordinates in standard space as independent variables:

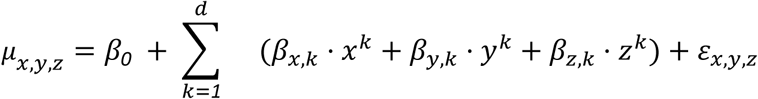

where *x*, *y* and *z* refer to mean-centered coordinates of one vertex inside the cluster and *d* = *5* is the order of polynomial expansion. This model was estimated using 10-fold cross-validation, predicting preferred numerosity in the left-out set of vertices. Predictive performance was assessed by calculating the correlation between actual and predicted preferred numerosity (cv-r) for which statistical significance can be determined (see Figure 6a,b and Supplementary Figure S7). Significant correlations between actual numerosities and predicted numerosities indicated that preferred numerosity changed along spatial dimensions and was thus topographically organized.

#### Statistical analysis of tuning parameters

Attempting to replicate previously reported effects^12^, we also analyzed relationships between preferred numerosity and cortical surface area (see Figure 5a,b and Supplementary Figure S5) and FWHM tuning width (see Figure 5c,d and Supplementary Figure S6) for each hemisphere. Additionally, we ran a linear mixed-effects model with either cortical surface area or FWHM tuning width as the dependent variable, preferred numerosity and hemisphere as independent variables and subject as a random effect (see Table 1).

#### Statistical analysis of behavioral performance

Bayesian correlations between the maximum cvR^2^ of the subjects and their behavioral performance (hit rates in the target detection task) were analyzed using JASP 0.16.4^43^. A uniform distribution on the range [-1,1] was chosen as the prior distribution.

## Data availability

After publication, the data for this study will be made available through a public link on the Edmond repository (https://edmond.mpg.de/). During the peer review phase, data are available from the corresponding author upon request.

## Code availability

All data analysis was performed with Spyder 5.4.3 using Python 3.10 and various third-party packages (numpy 1.24.3, scipy 1.10.1, pandas 2.1.1, statsmodels 0.14.0, nibabel 5.1.0), except for edge-based surface clustering which was run using AFNI’s SurfClust. Brain surfaces were visualized with surfplot 0.2.0 and other plots were generated with matplotlib 3.7.1. All code for data analysis is available from GitHub (https://github.com/SkeideLab/EMPRISE-KIDS).

## Acknowledgements

We thank our lab manager Micha Vollmann for recruiting participants, coordinating logistics, handling forms, and organizing the measurements. This work was supported by the German Research Foundation (DFG Research Grant 433715509 and DFG Heisenberg Program Grant 433758790 awarded to M.A.S.) and the Jacobs Foundation (Research Fellowship awarded to M.A.S.).

## Author contributions

M.A.S. conceived the study. R.T. performed experiments. G.J. and J.S. analyzed data. G.J. and J.S. visualized the results. M.A.S. wrote the manuscript with feedback from all other authors. M.A.S. supervised the study.

## Ethics declarations

The authors declare no competing interests.

