## Supplementary Material for "High-field fMRI at 7 Tesla reveals topographic responses tuned to number in the developing human brain"

[1] Research Group Learning in Early Childhood,
Max Planck Institute for Human Cognitive and Brain Sciences,
Stephanstraße 1A, 04103 Leipzig, Germany

[2] Institute for Psychology,
Otto-von-Guericke-Universität Magdeburg
Universitätsplatz 2, 39106 Magdeburg, Germany

[3] Department of Neurophysics,
Max Planck Institute for Human Cognitive and Brain Sciences,
Stephanstraße 1A, 04103 Leipzig, Germany

[4] Animal Physiology Unit, Institute of Neurobiology,
Eberhard-Karls-Universität Tübingen,
Auf der Morgenstelle 28, 72076 Tübingen, Germany

[5] These authors contributed equally: Ga-Ram Jeong, Joram Soch.

✉

Correspondence should be addressed to Michael A. Skeide

### Supplementary Tables

| subject | gender | age | hit rate | max cvR² | mean cvR² |
|---|---|---|---|---|---|
| c001 | F | 12 | 98.84 | 0.57 | 0.27 |
| c002 | M | 12 | 99.42 | 0.44 | 0.22 |
| c003 | F | 12 | 99.71 | 0.41 | 0.22 |
| c004 | M | 12 | 95.93 | 0.49 | 0.23 |
| c005 | F | 12 | 100.00 | 0.69 | 0.34 |
| c006 | M | 12 | 98.83 | 0.66 | 0.34 |
| c007 | M | 12 | 89.24 | 0.49 | 0.24 |
| c008 | M | 12 | 98.55 | 0.60 | 0.28 |
| c009 | M | 12 | 98.55 | 0.55 | 0.26 |
| c010 | F | 12 | 99.71 | 0.73 | 0.37 |
| c011 | M | 12 | 98.53 | 0.69 | 0.34 |
| c012 | M | 11 | 93.90 | 0.44 | 0.24 |
| a001 | F | 27 | 94.74 | 0.63 | 0.30 |
| a002 | M | 26 | 79.46 | 0.42 | 0.23 |
| a003 | M | 22 | 100.00 | 0.72 | 0.38 |
| a004 | F | 30 | 99.42 | 0.71 | 0.35 |
| a005 | F | 24 | 98.25 | 0.61 | 0.30 |
| a006 | M | 20 | 96.77 | 0.47 | 0.24 |
| a007 | F | 20 | 94.17 | 0.54 | 0.29 |
| a008 | M | 27 | 78.31 | 0.37 | 0.20 |
| a009 | F | 22 | 98.24 | 0.65 | 0.32 |
| a010 | M | 22 | 99.42 | 0.70 | 0.36 |
| a011 | F | 34 | 99.13 | 0.67 | 0.32 |
| a012 | M | 33 | 90.12 | 0.49 | 0.23 |

**Table S1 | Participant details and behavioral task performance.** The table shows the pseudonymous subject ID, gender (F = female, M = male), age in years, catch trial hit rates (averaged across runs), maximum and mean variance explained by the model (maximum and mean of supra-threshold $cvR^2$) averaged across both hemispheres.

| Figure | tuning parameters | native coordinates | standard coordinates | anatomical location (AAL regions) |
|---|---|---|---|---|
| 2a/b | mu = 1.60, fwhm = 3.48 | [-15, 15, 53] | [-22, -3, 50 ] | Frontal_Sup_L (NF) |
| 2c/d | mu = 4.05, fwhm = 16.7 | [-25, -61, 11] | [-47,-71, -6 ] | Occipital_Inf_L (NTO) |
| 2e/f | mu = 1.80, fwhm = 3.31 | [-18, -26, 55] | [-29, -52, 46] | Parietal_Inf_L (NPC1) |
| 2g/h | mu = 4.40, fwhm = 19.4 | [-46, -55, 7] | [-44, -83, -4] | Occipital_Inf_L (NTO) |

**Table S2 | Anatomical locations.** Automated anatomical labeling (AAL) regions for vertices reported in Figure 2. mu = preferred numerosity; fwhm = full width at half maximum. Sup = superior. Inf = inferior. L = left. NTO = temporo-occipital visual numerosity field; NPC1 = parietal visual numerosity fields; NF = frontal visual numerosity field.

### Supplementary Figures

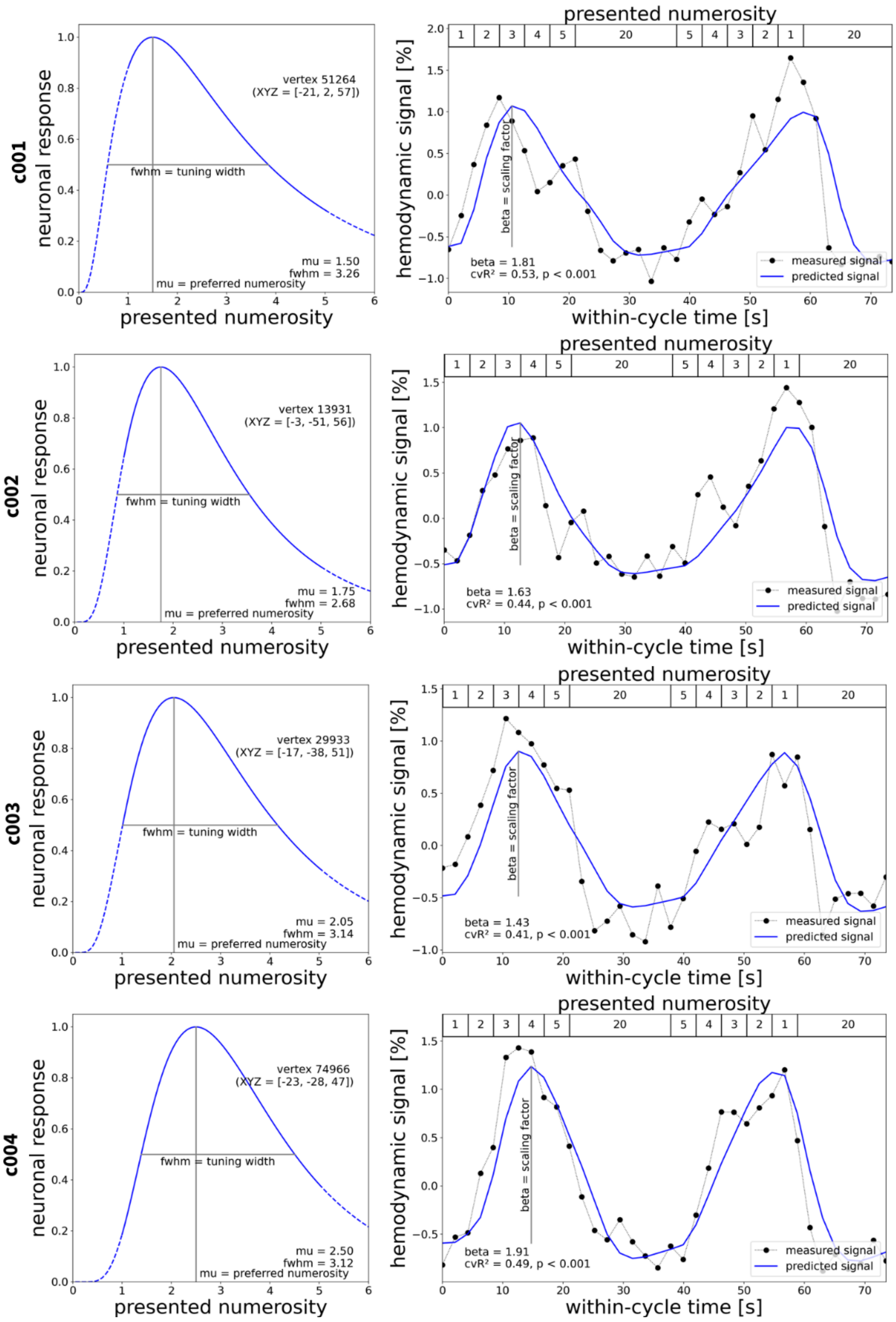


**Figure S1 | Neural tuning and hemodynamic responses in children (page 1 of 3).** Neural response functions (left) and hemodynamic responses measured using fMRI during visual numerosity presentation (right) were extracted from the vertex with the highest $cvR^2$.

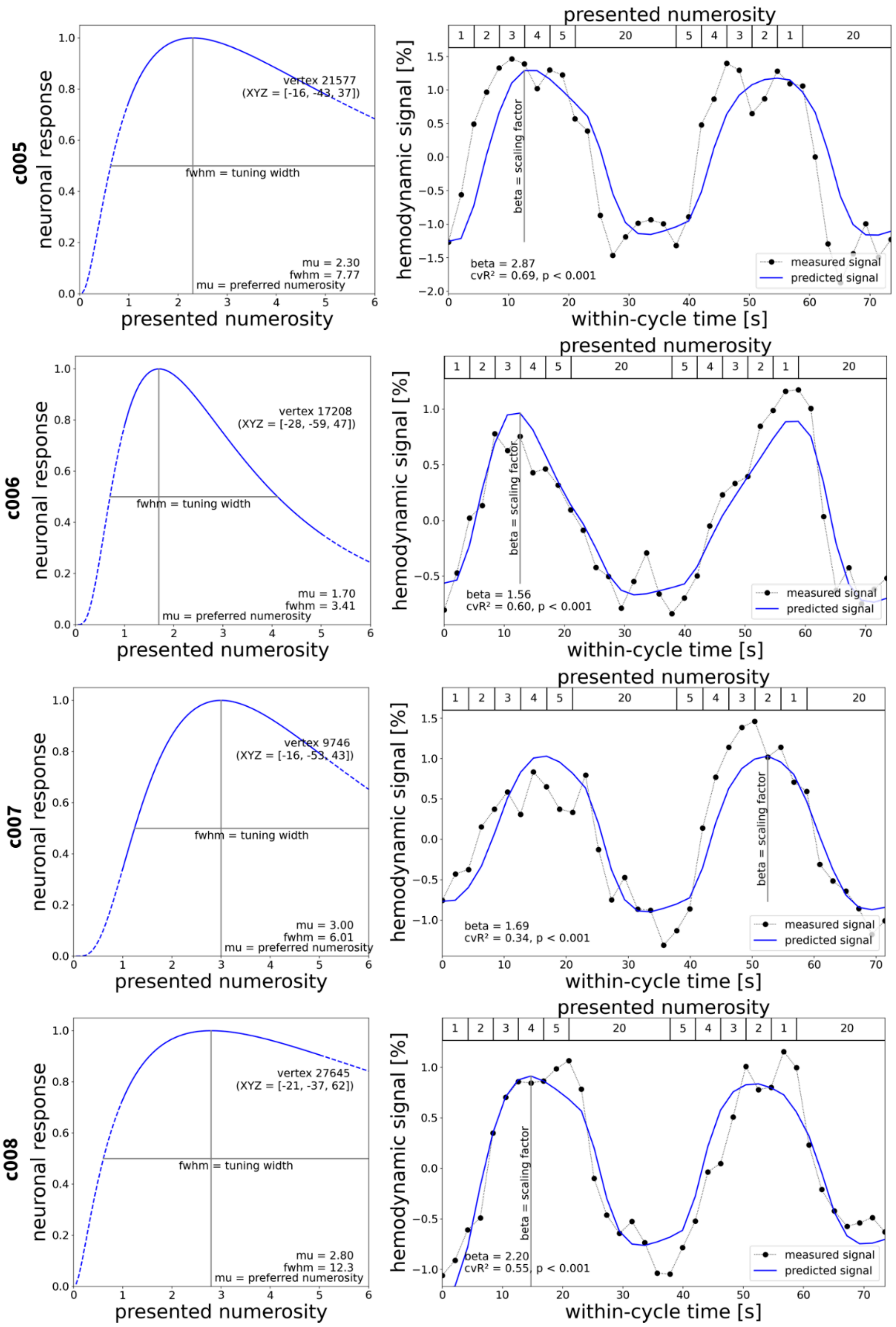


**Figure S1 | Neural tuning and hemodynamic responses in children (page 2 of 3).**

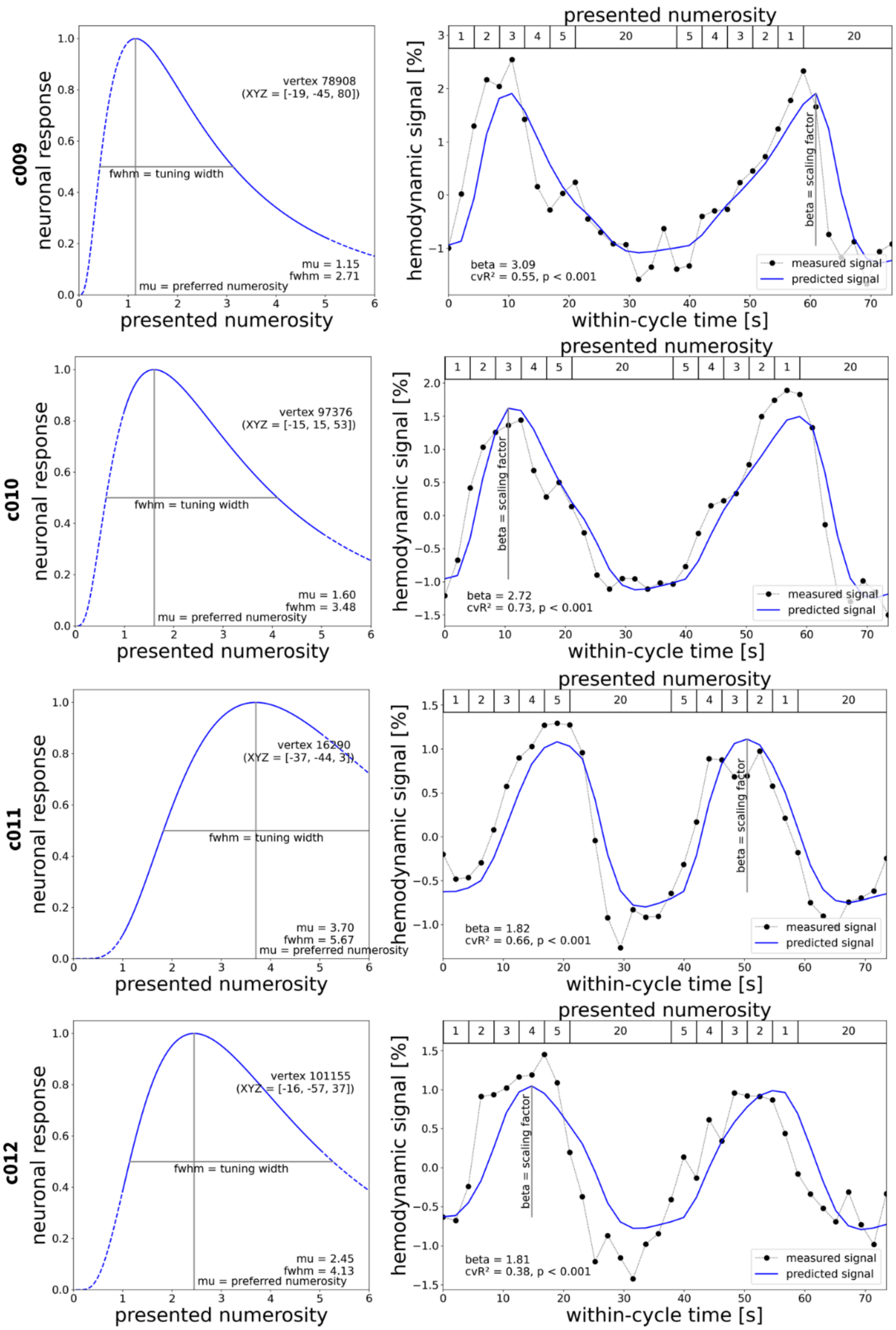


**Figure S1 | Neural tuning and hemodynamic responses in children (page 3 of 3).**

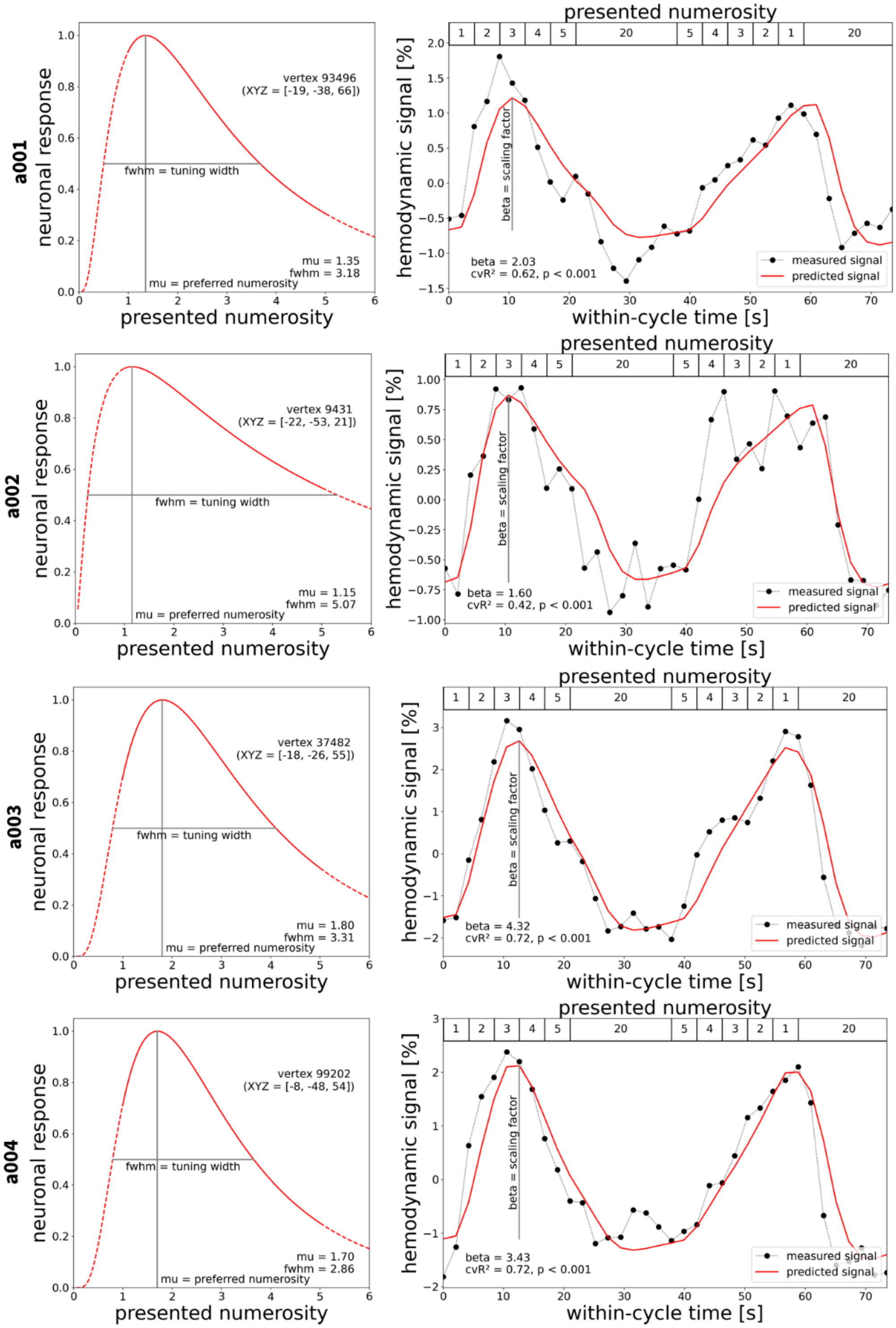


**Figure S2 | Neural tuning and hemodynamic responses in adults (page 1 of 3).** Neural response functions (left) and hemodynamic responses measured using fMRI during visual numerosity presentation (right) were extracted from the vertex with the highest $cvR^2$.

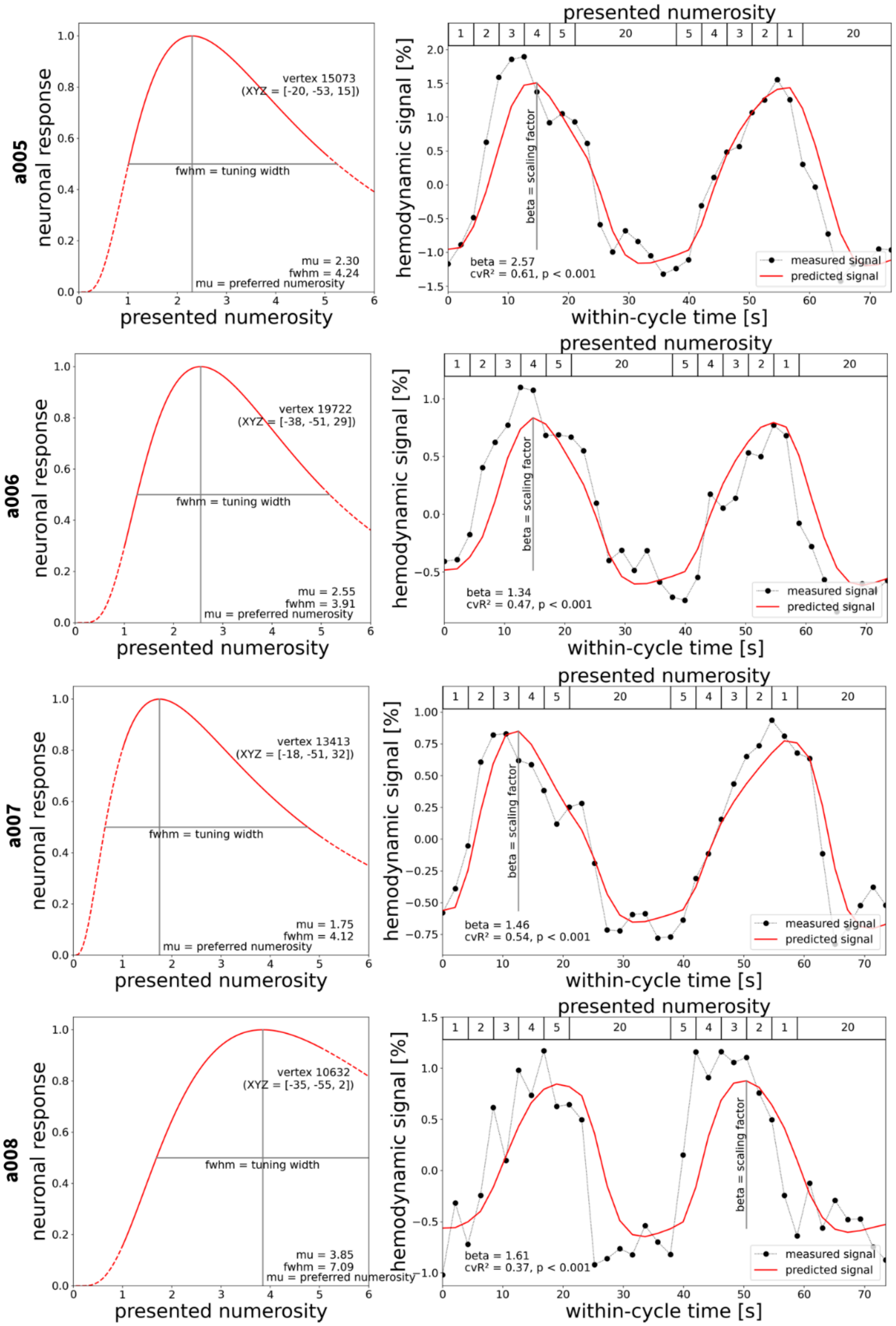


**Figure S2 | Neural tuning and hemodynamic responses in adults (page 2 of 3).**

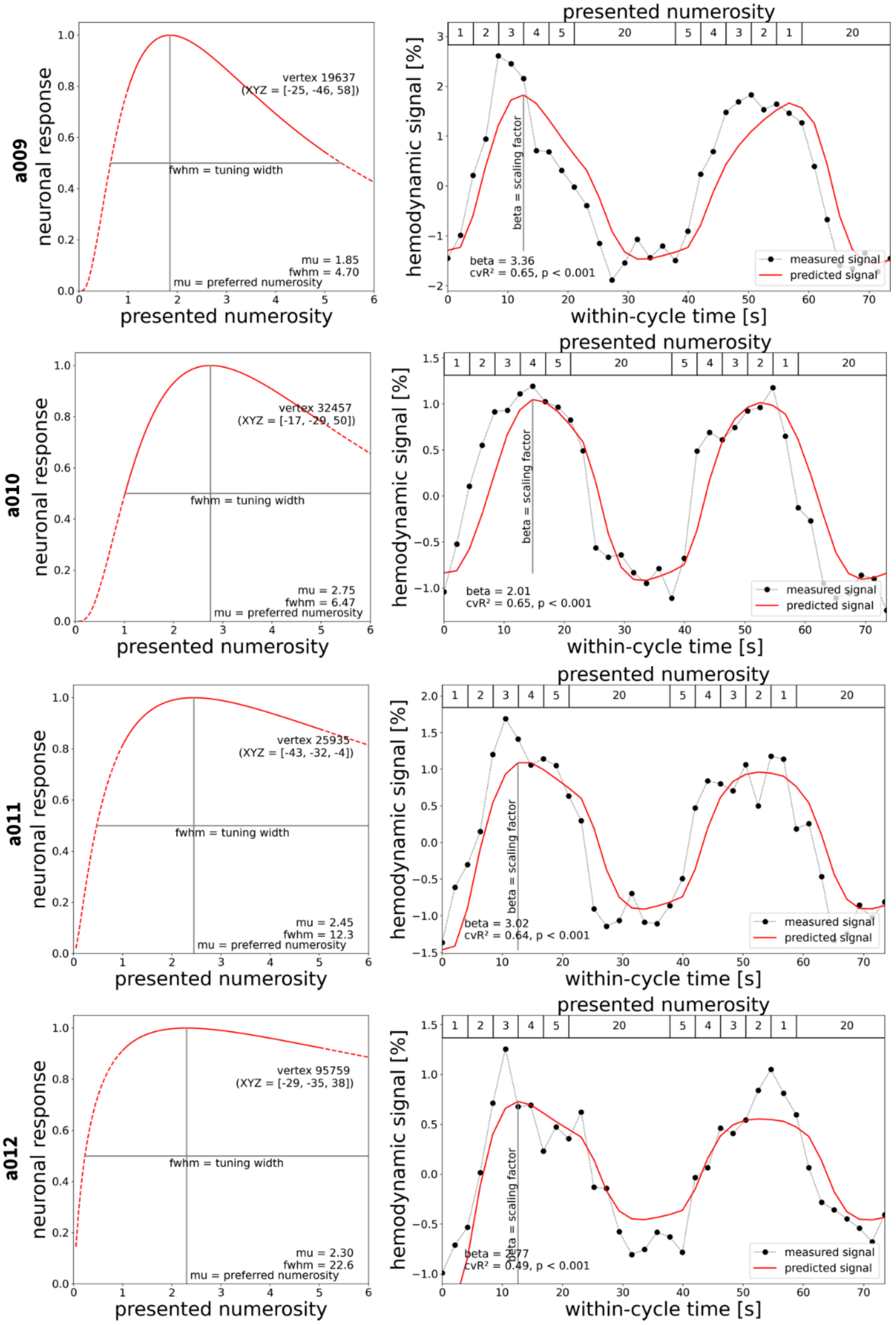


**Figure S2 | Neural tuning and hemodynamic responses in adults (page 3 of 3).**

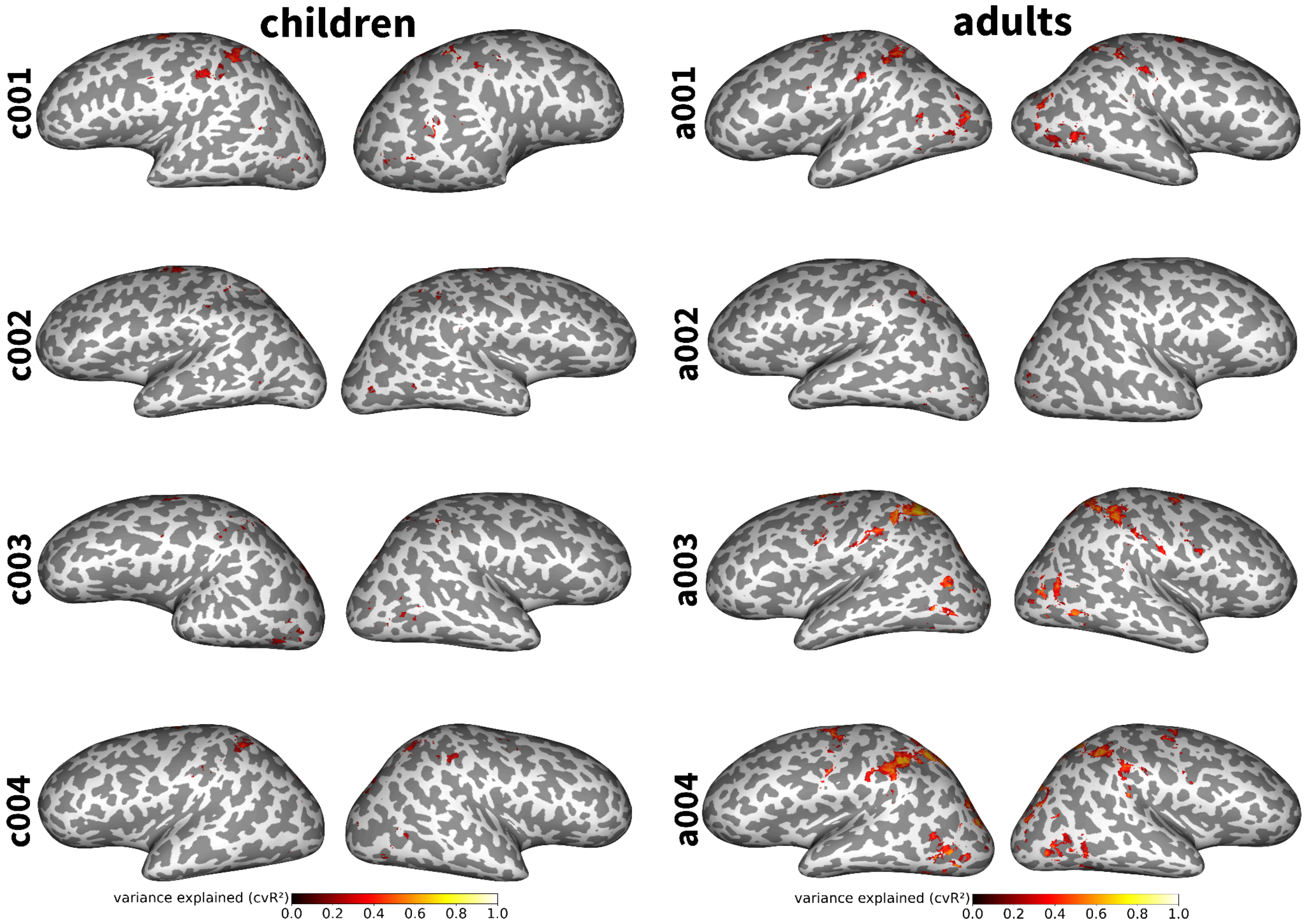


**Figure S3 | Variance explained by the neural tuning model (page 1 of 3).** Inflated surface maps show $R^2$ of vertices in each subject's native space in which the neural tuning model explains a significant amount of the variance ($p < 0.05$, Bonferroni-corrected) during visual numerosity perception.

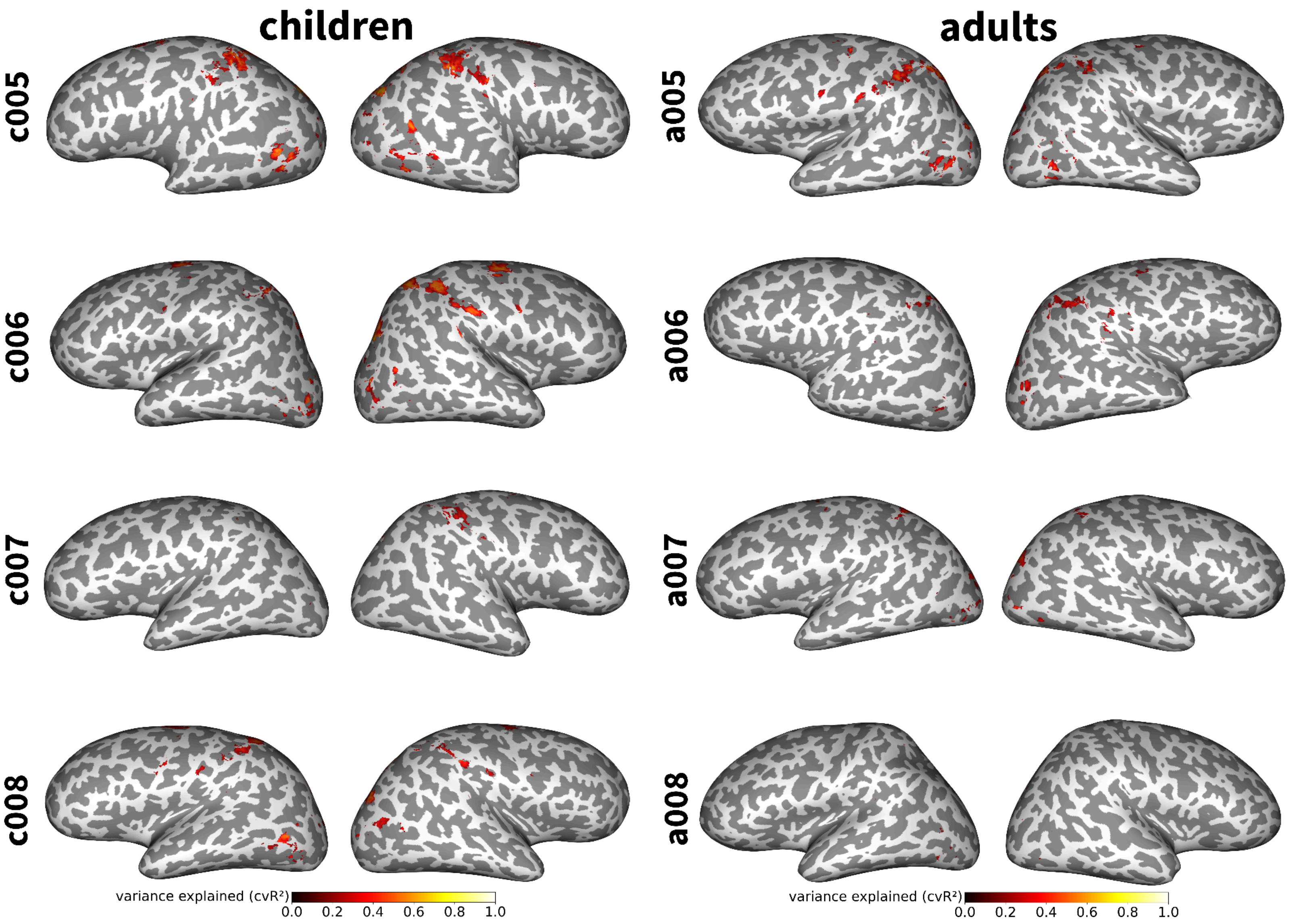


**Figure S3 | Variance explained by the neural tuning model (page 2 of 3).**

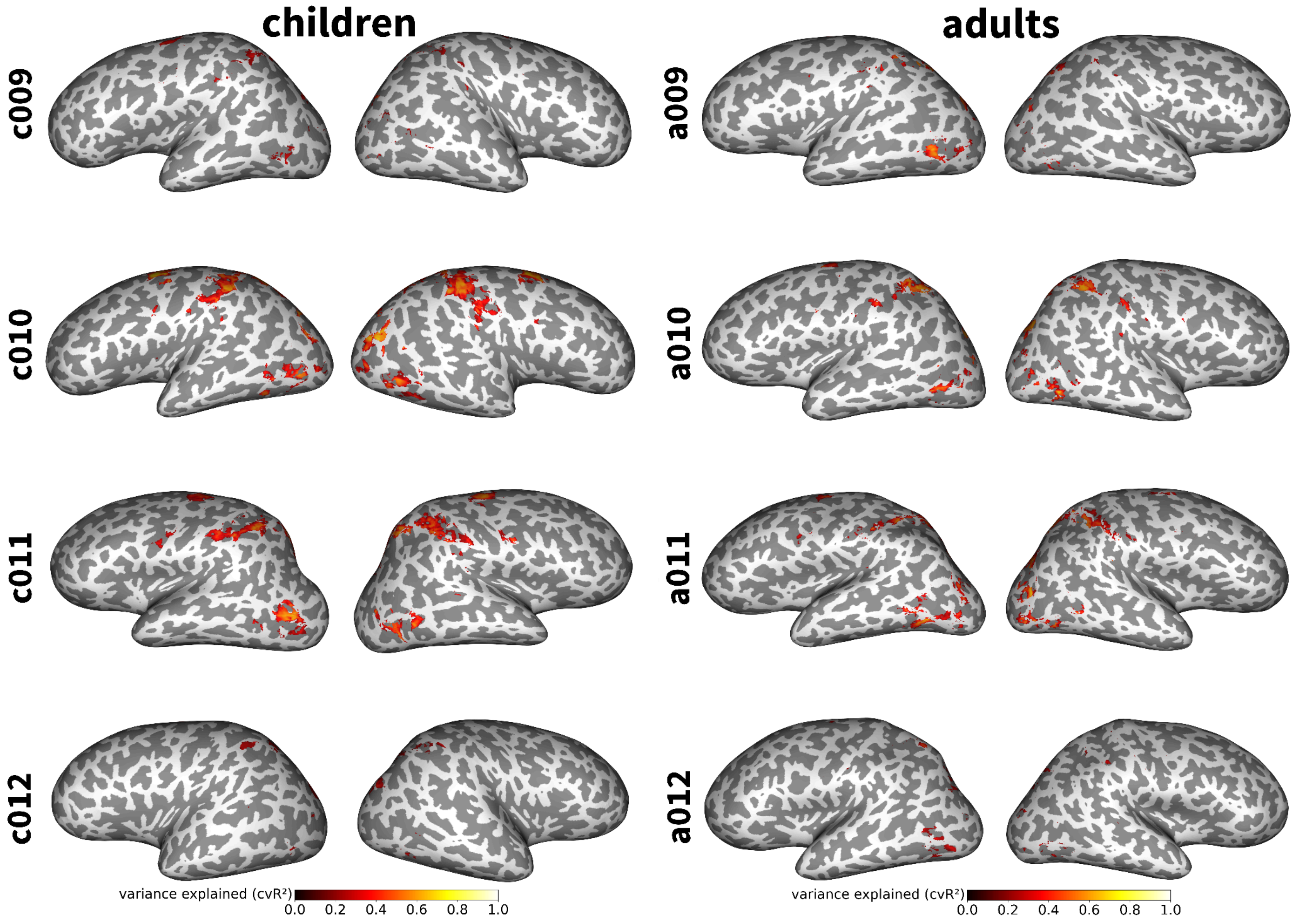


**Figure S3 | Variance explained by the neural tuning model (page 3 of 3).**

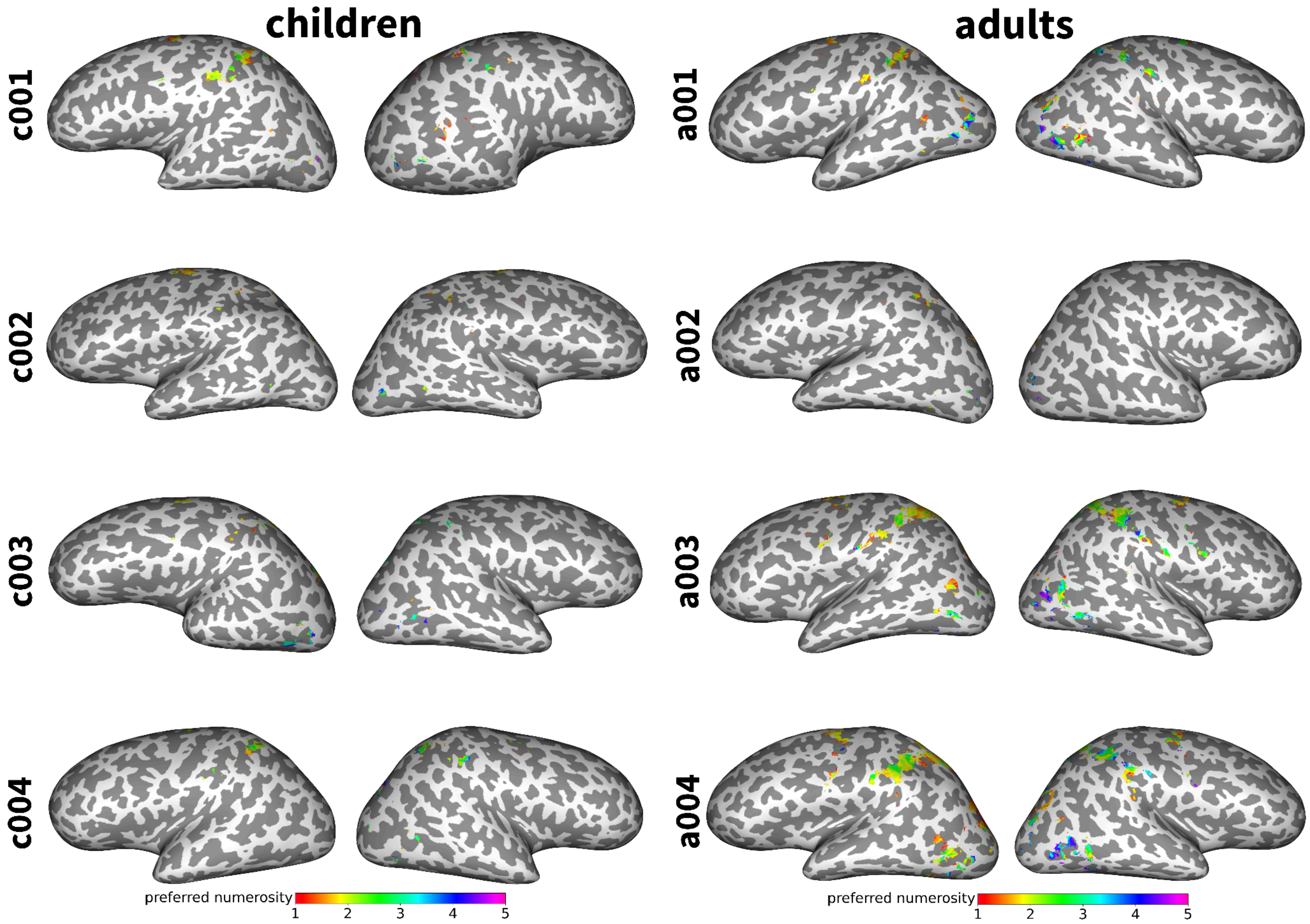


**Figure S4 | Preferred numerosity maps (page 1 of 3).** Inflated surface maps show preferred numerosity of vertices in each subject's native space in which the neural tuning model explains a significant amount of the variance during ($p < 0.05$, Bonferroni-corrected) visual numerosity perception.

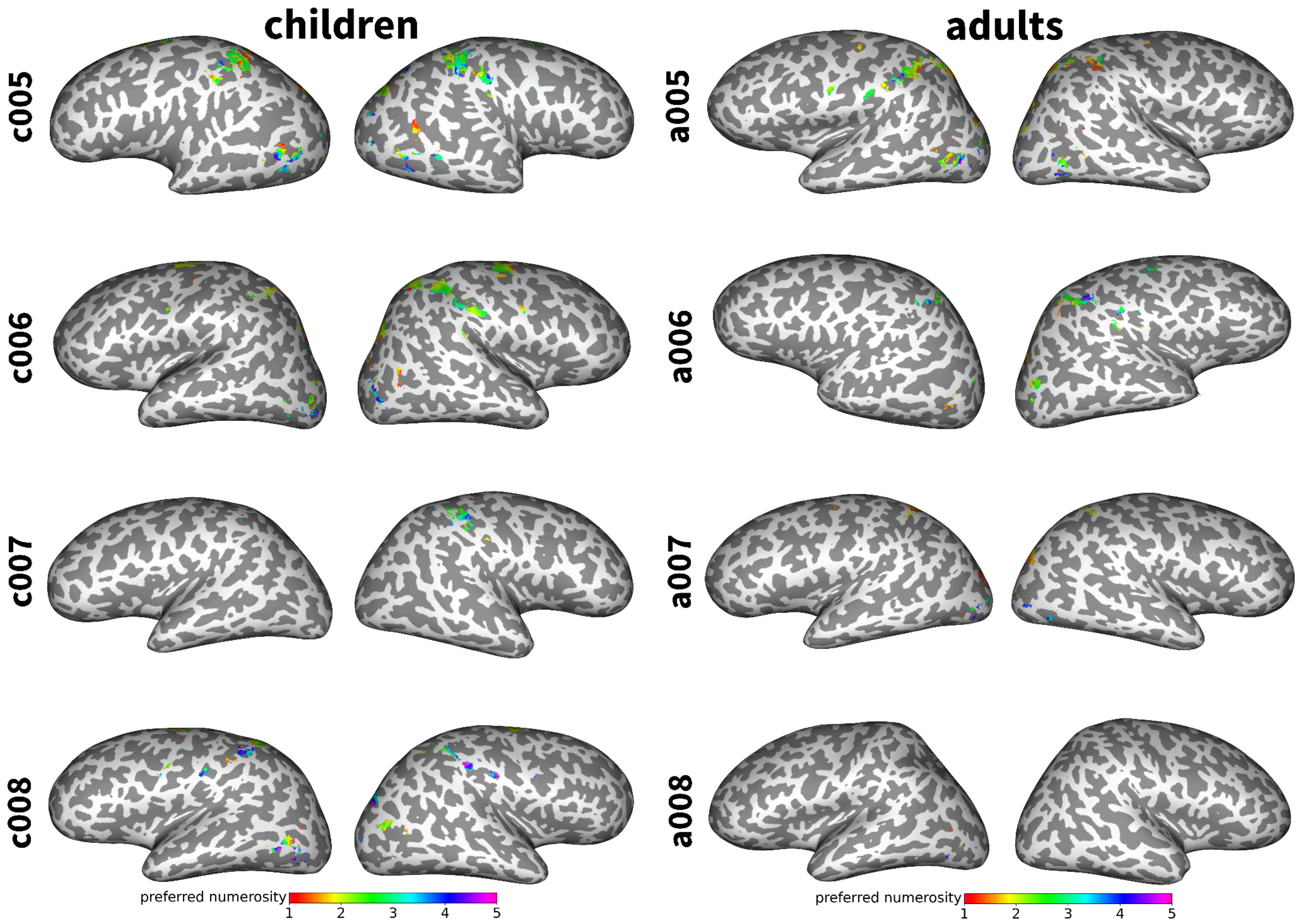


**Figure S4 | Preferred numerosity maps (page 2 of 3).**

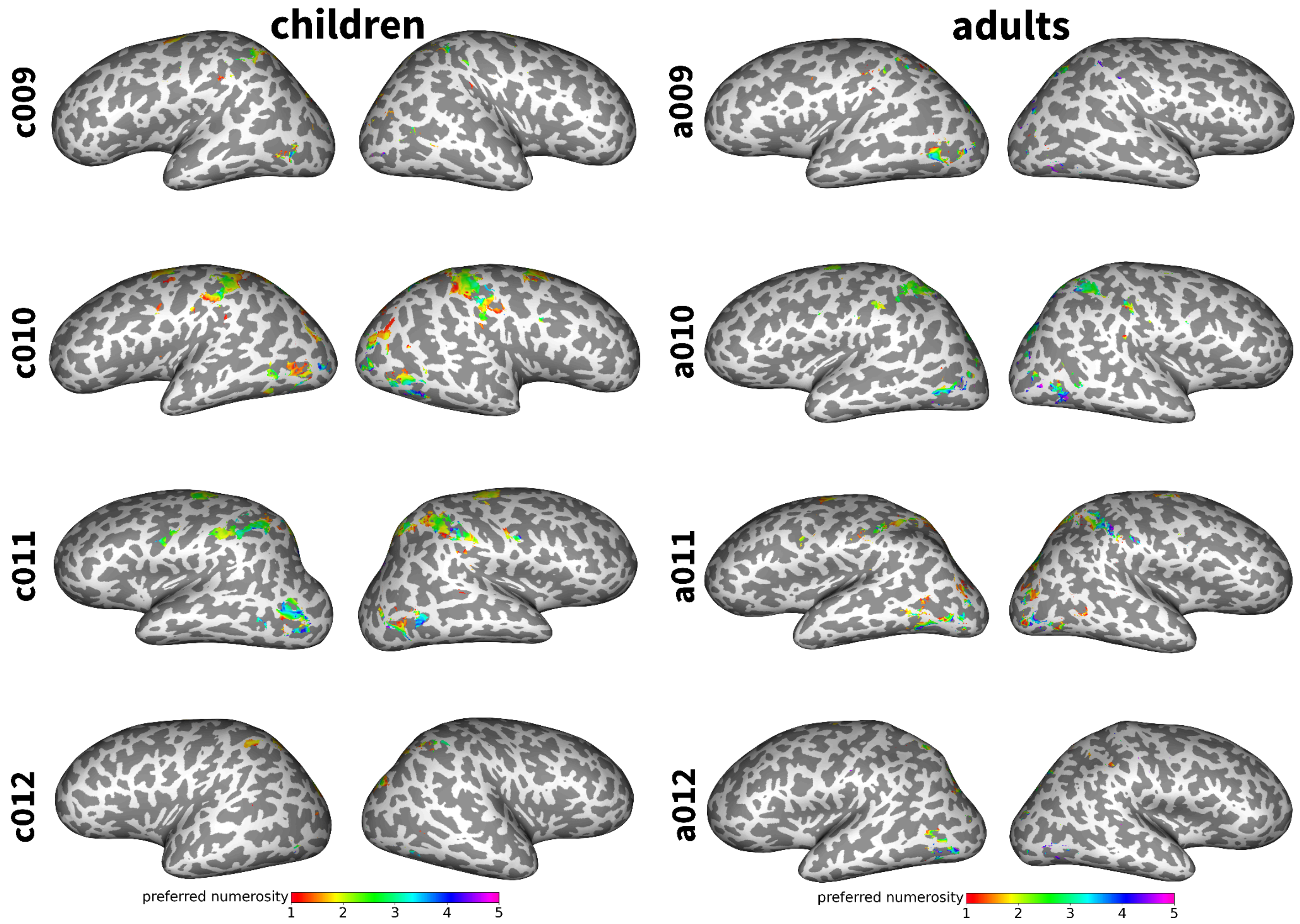


**Figure S4 | Preferred numerosity maps (page 3 of 3).**

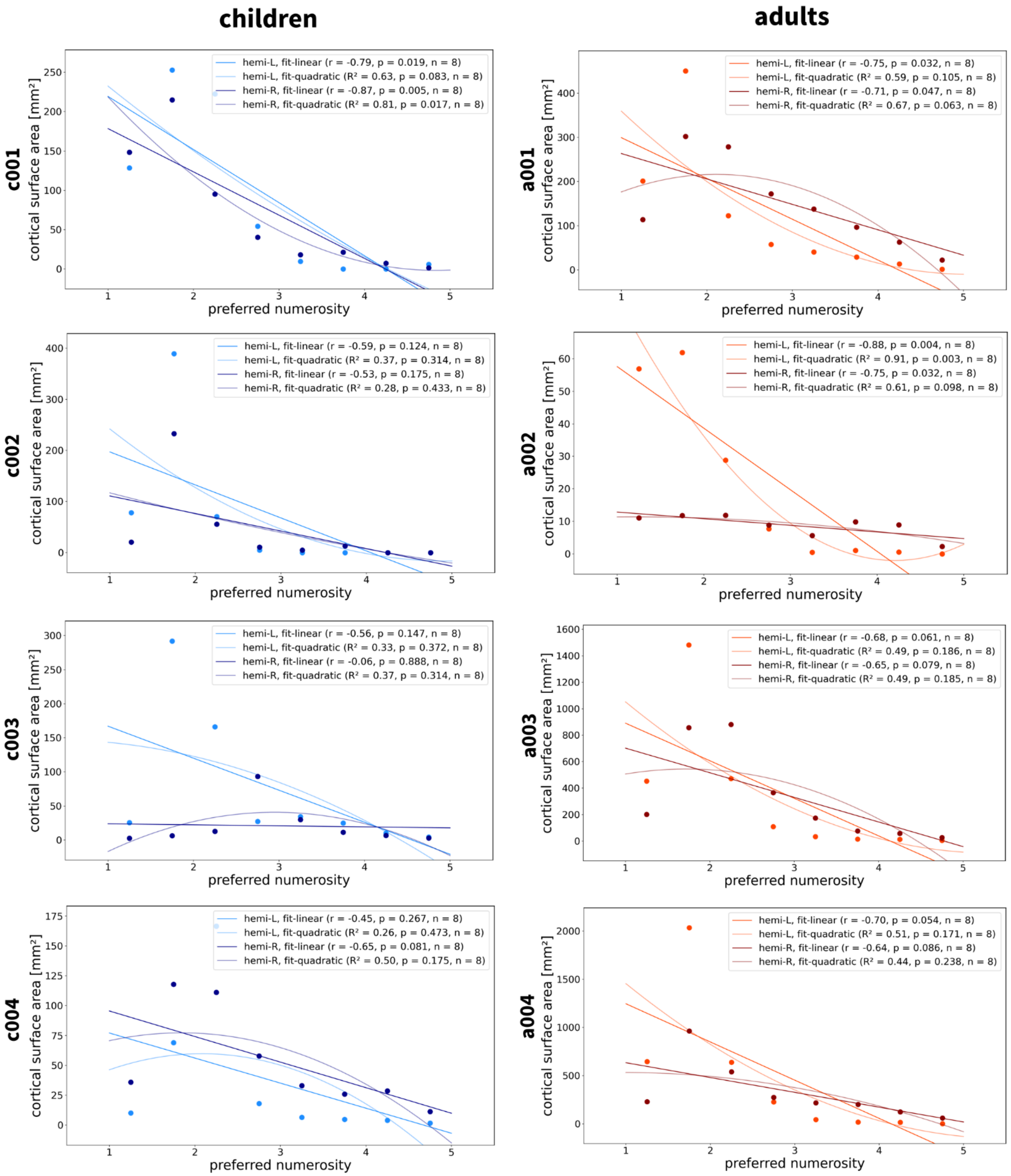


**Figure S5 | Associations between preferred numerosity and surface area (page 1 of 3).** Cortical surface area decreases as a function of preferred numerosity in most children (left) and adults (right).

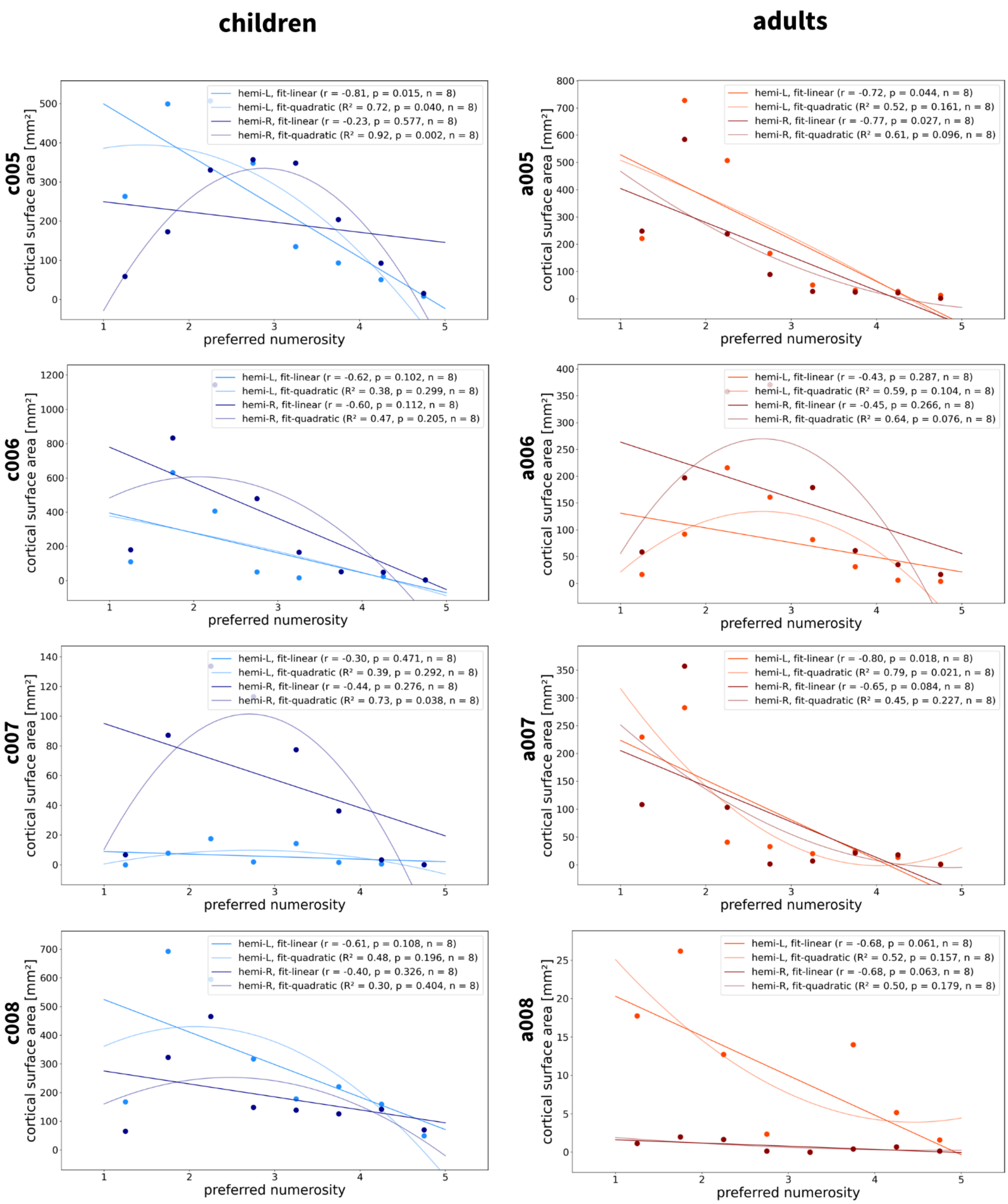


**Figure S5 | Associations between preferred numerosity and surface area (page 2 of 3).**

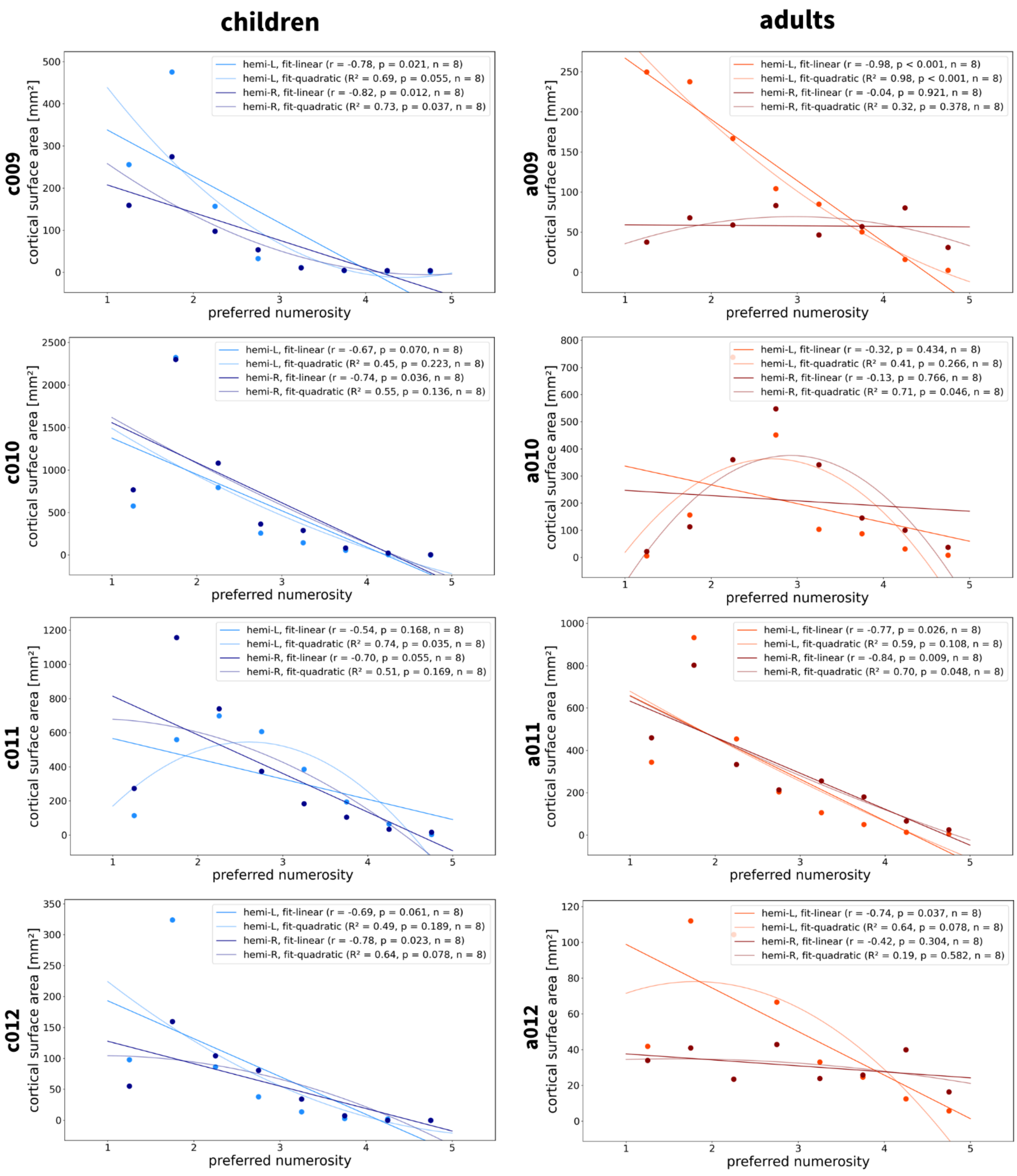


**Figure S5 | Associations between preferred numerosity and surface area (page 3 of 3).**

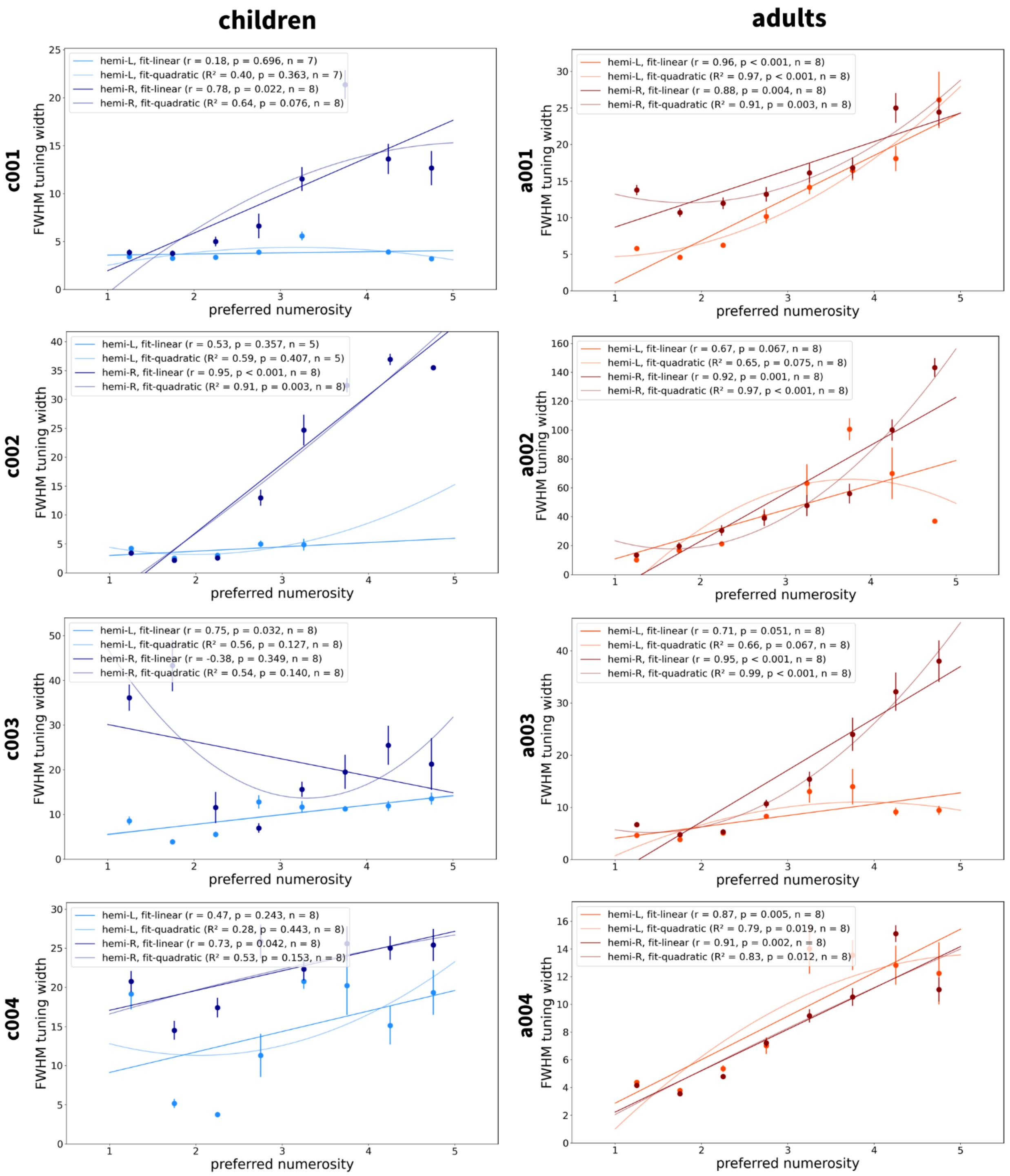


**Figure S6 | Associations between preferred numerosity and tuning width (page 1 of 3).** Full width at half maximum (FWHM) increases as a function of preferred numerosity in most children (left) and adults (right).

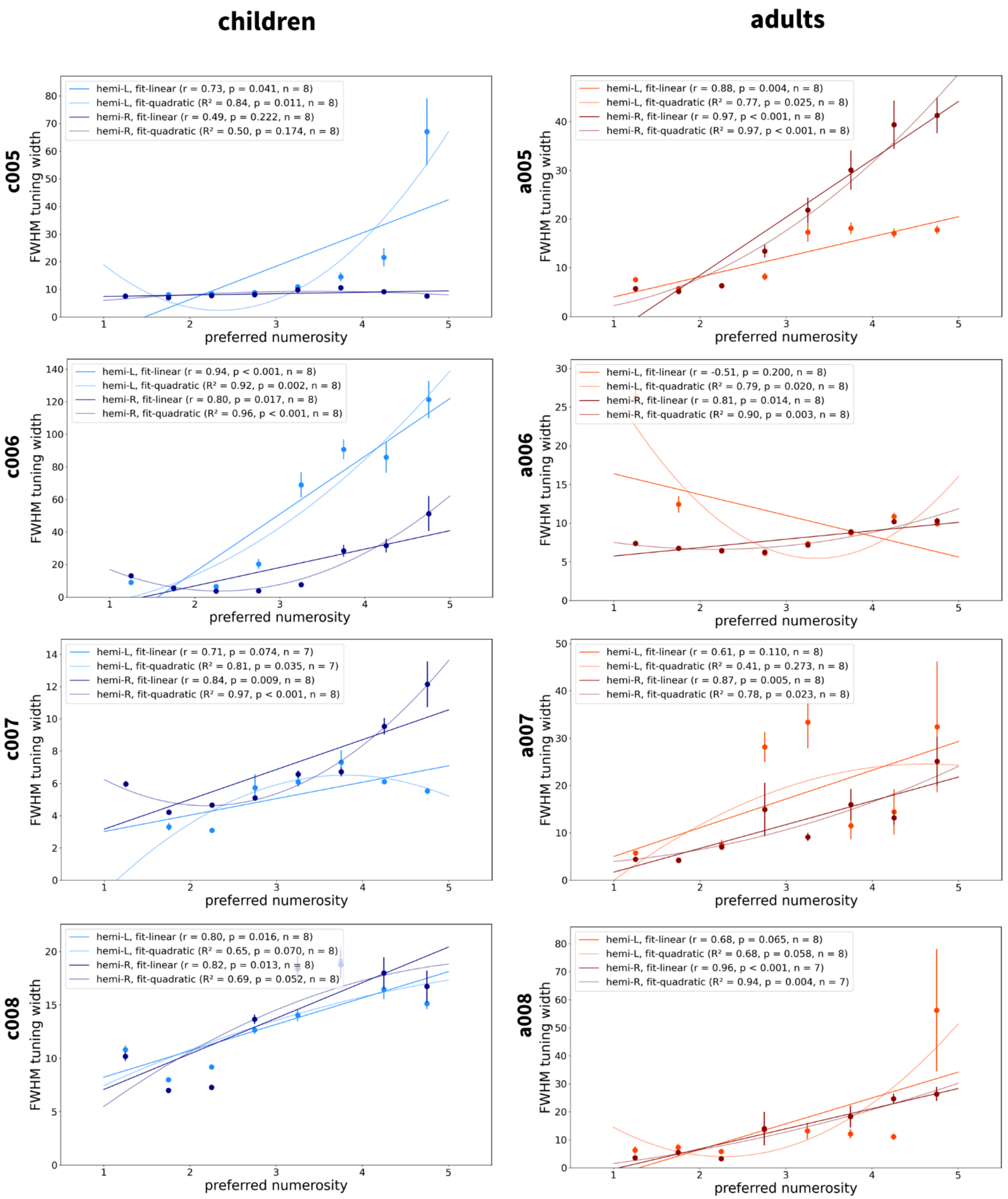


**Figure S6 | Associations between preferred numerosity and tuning width (page 2 of 3).**

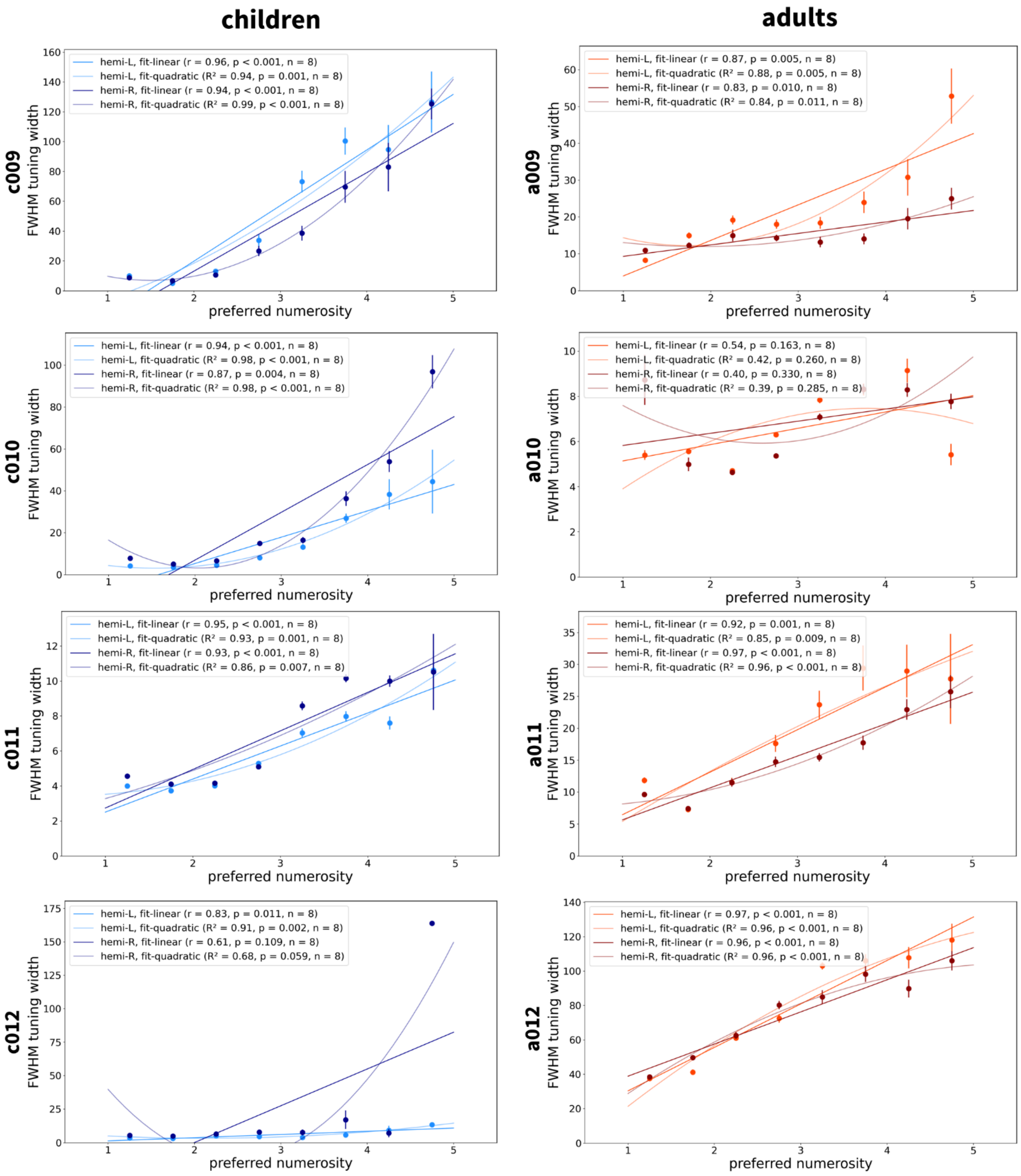


**Figure S6 | Associations between preferred numerosity and tuning width (page 3 of 3).**

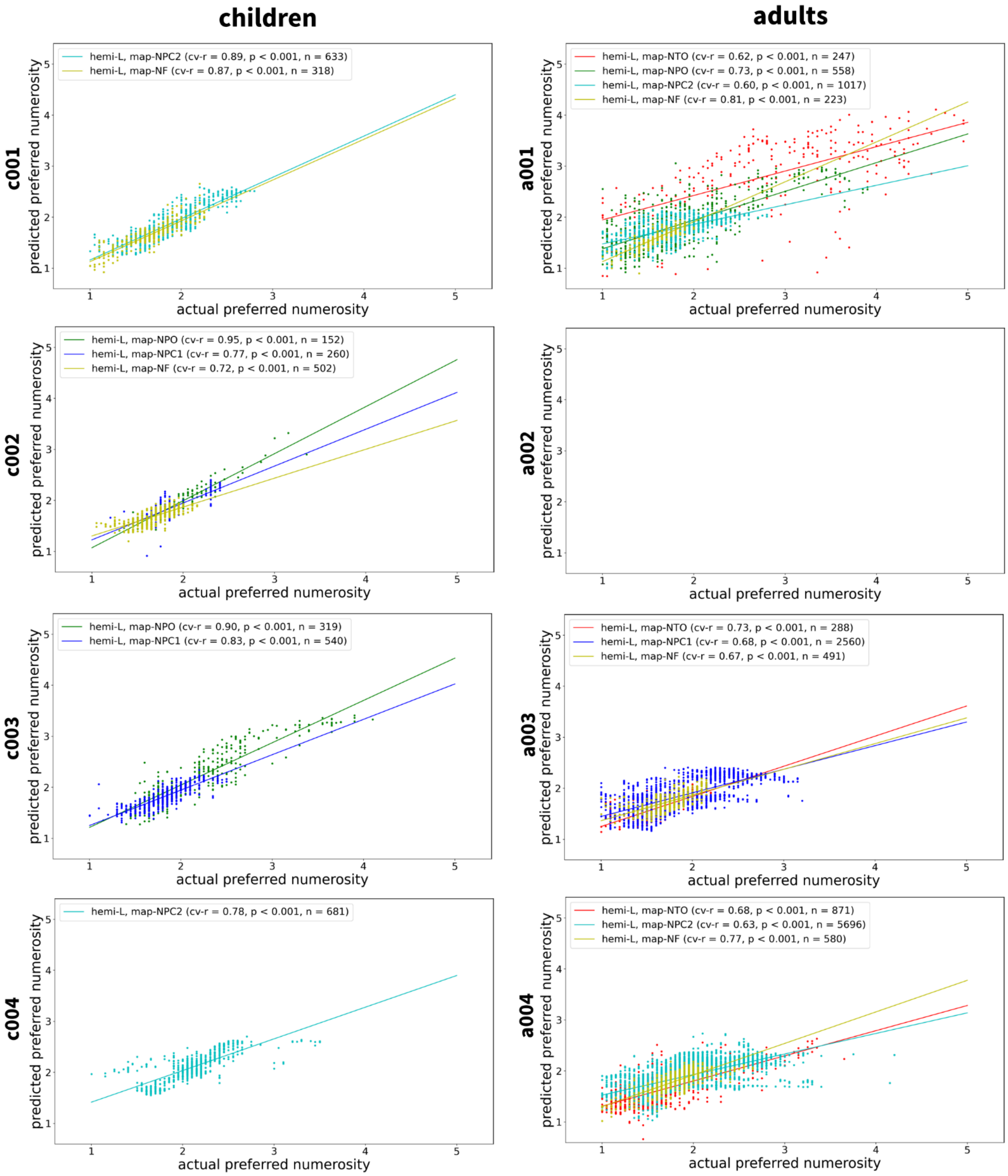


**Figure S7 | Fitted and preferred numerosity of topographic maps (page 1 of 3).** Preferred numerosity was predicted from surface coordinates, indicating a progression of preferred numerosity on the cortical surface within topographic maps. No data are shown in accordance with predefined thresholds whenever clusters did not exceed a surface area of 50 $mm^2$ or when the minimum distance of a cluster from prespecified numerosity maps was larger than 25 mm.

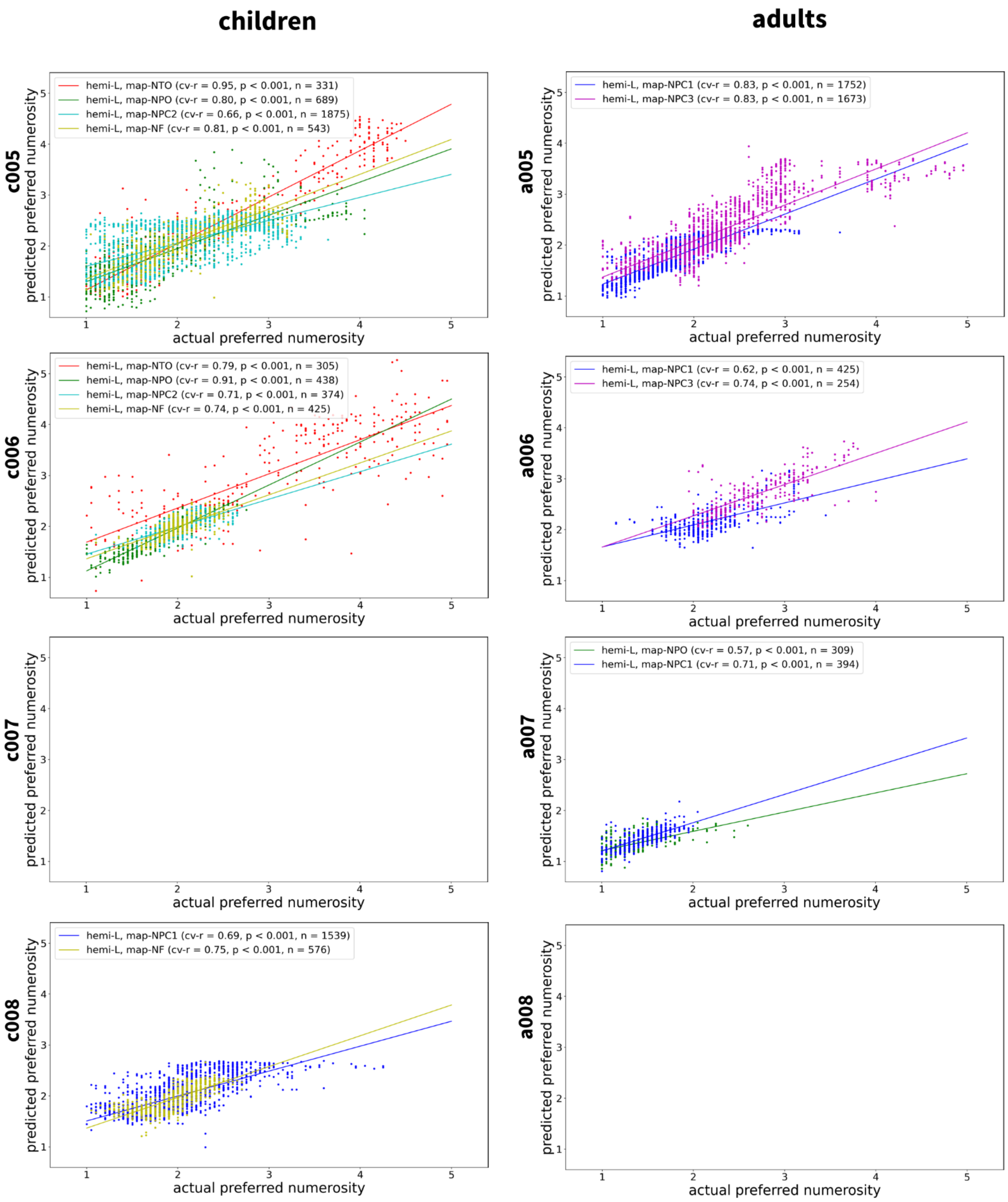


**Figure S7 | Fitted and preferred numerosity of topographic maps (page 2 of 3).**

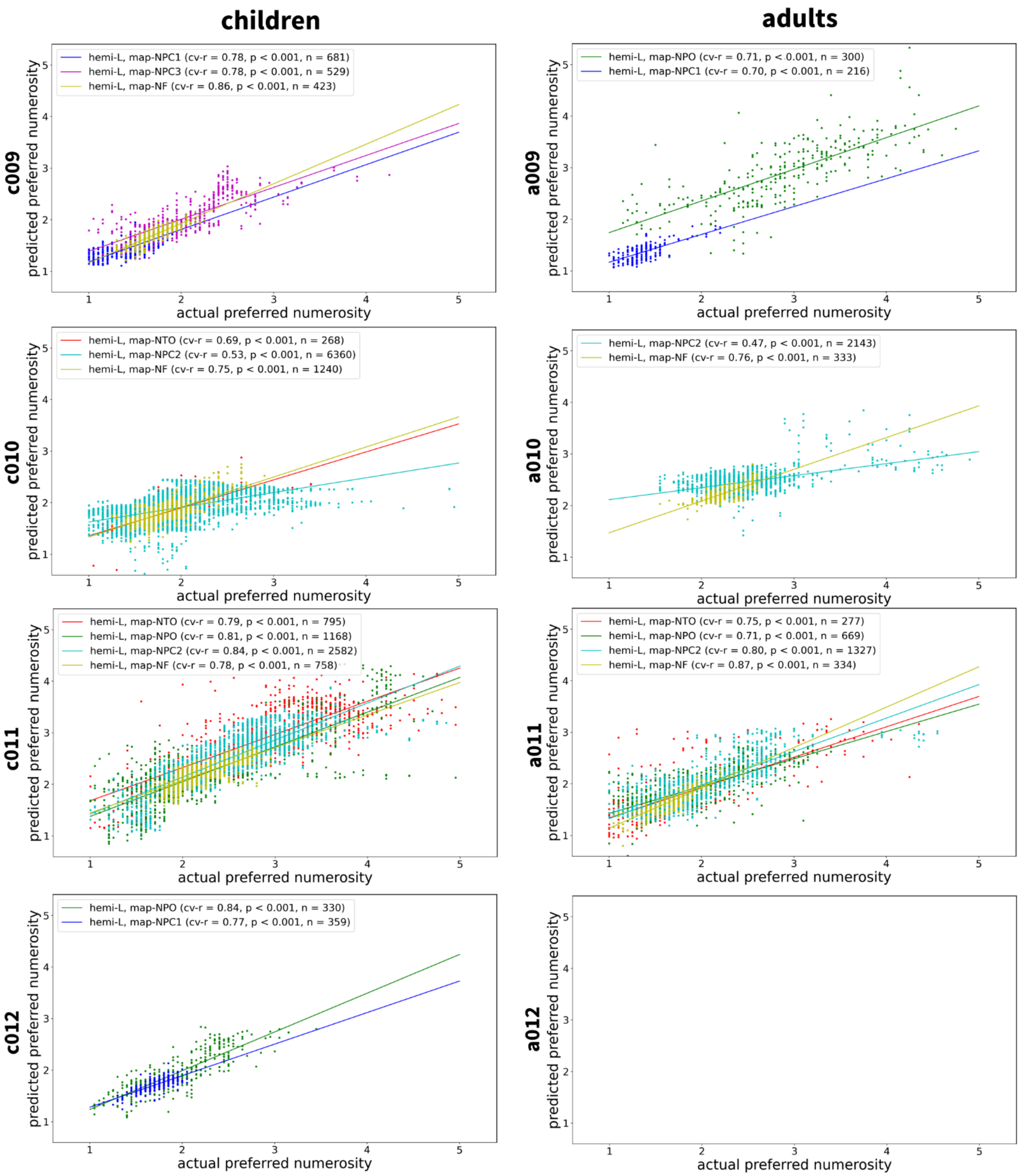


**Figure S7 | Fitted and preferred numerosity of topographic maps (page 3 of 3).**

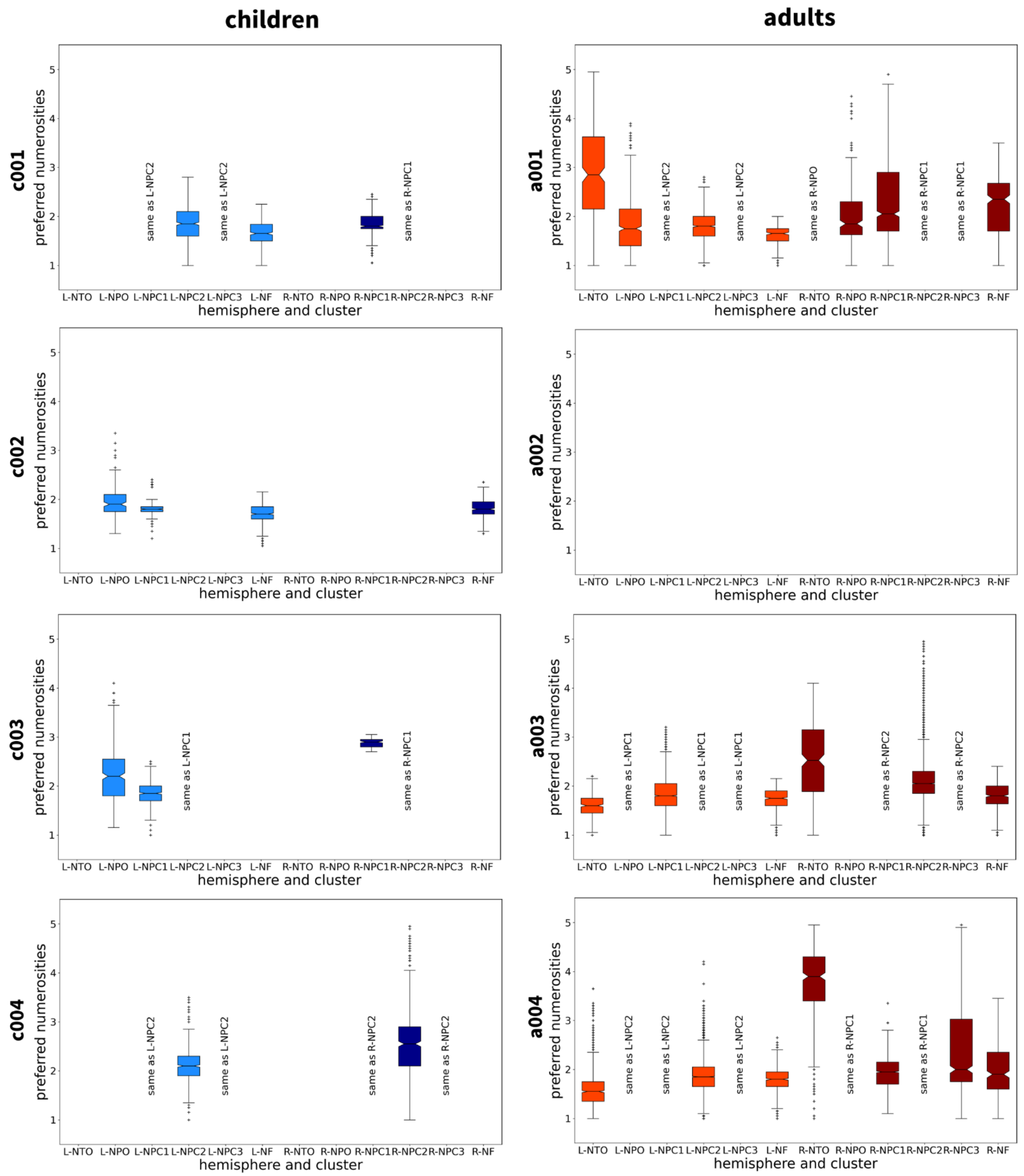


**Figure S8 | Range of preferred numerosities within topographic maps (page 1 of 3).** Preferred numerosities of all supra-threshold vertices within up to six numerosity maps per hemisphere are shown as boxplots. Boxes depict the interquartile range (IQR). Horizontal lines within boxes mark the median. Whiskers indicate the most extreme points that are still within 1.5 x IQR from the upper or lower quartile. Crosses denote data points outside this range. No data are shown in accordance with predefined thresholds whenever clusters did not exceed a surface area of 50 mm$^2$ or when the minimum distance of a cluster from prespecified numerosity maps was larger than 25 mm.

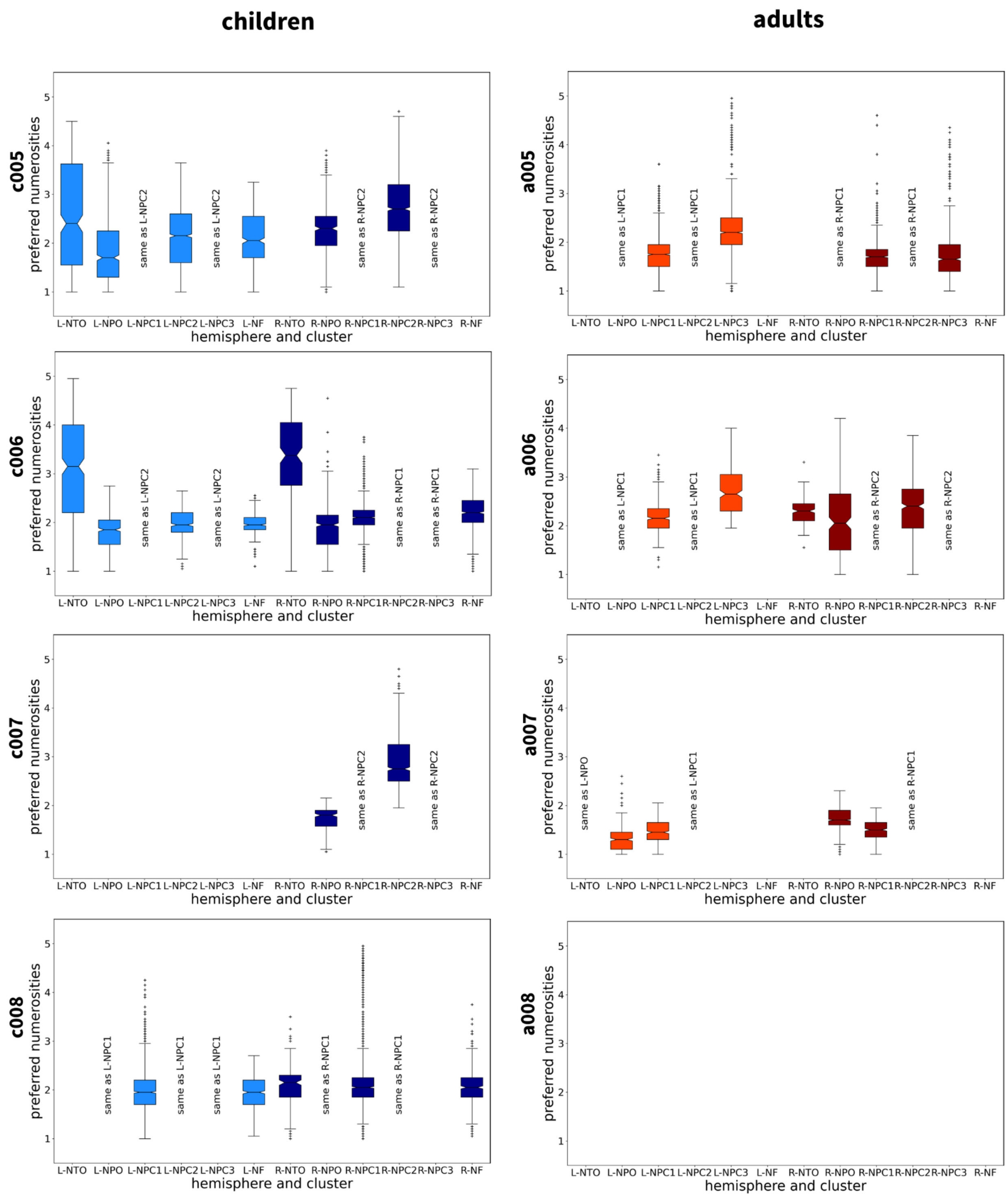


**Figure S8 | Range of preferred numerosities within topographic maps (page 2 of 3).**

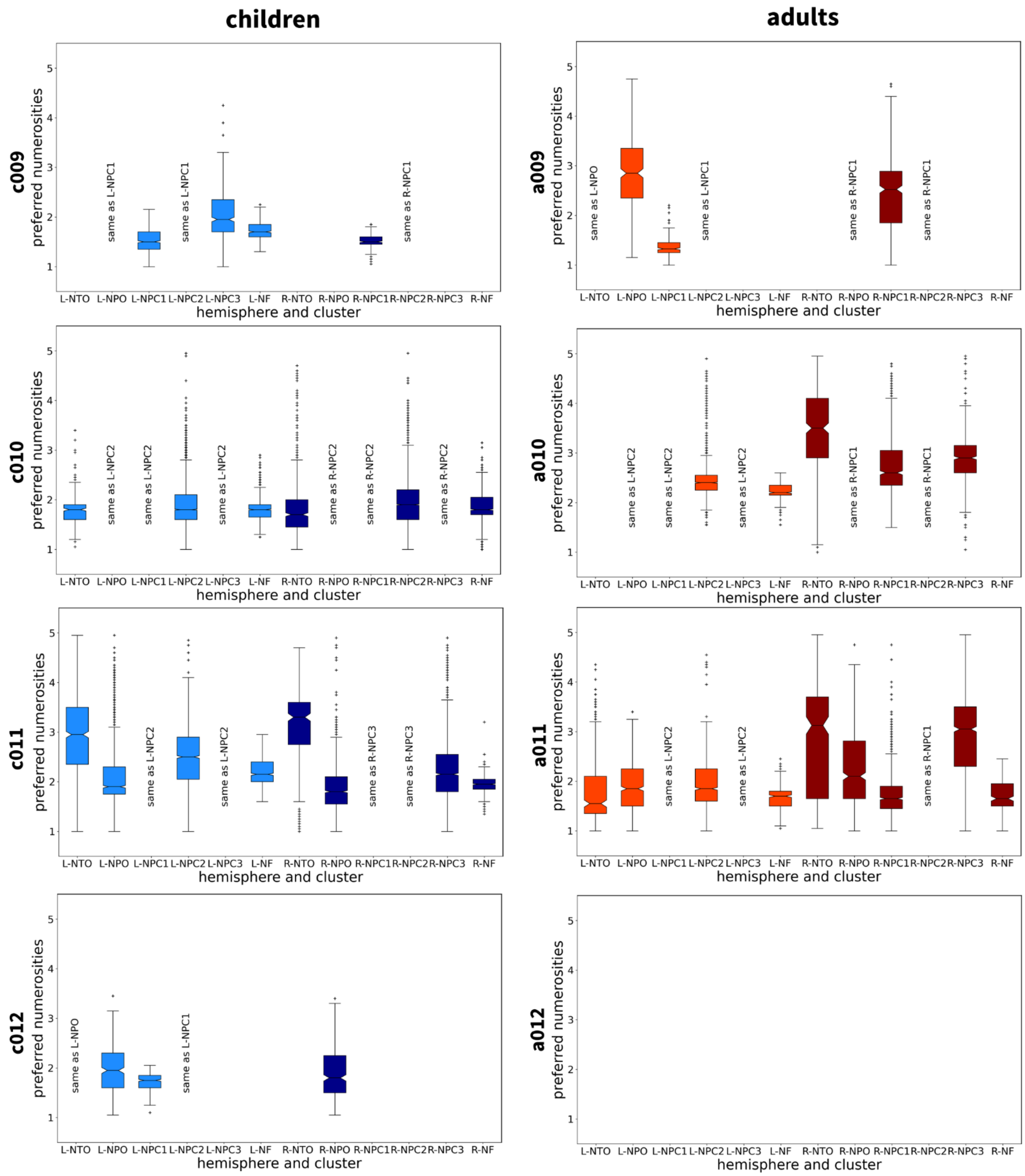


**Figure S8 | Range of preferred numerosities within topographic maps (page 3 of 3).**

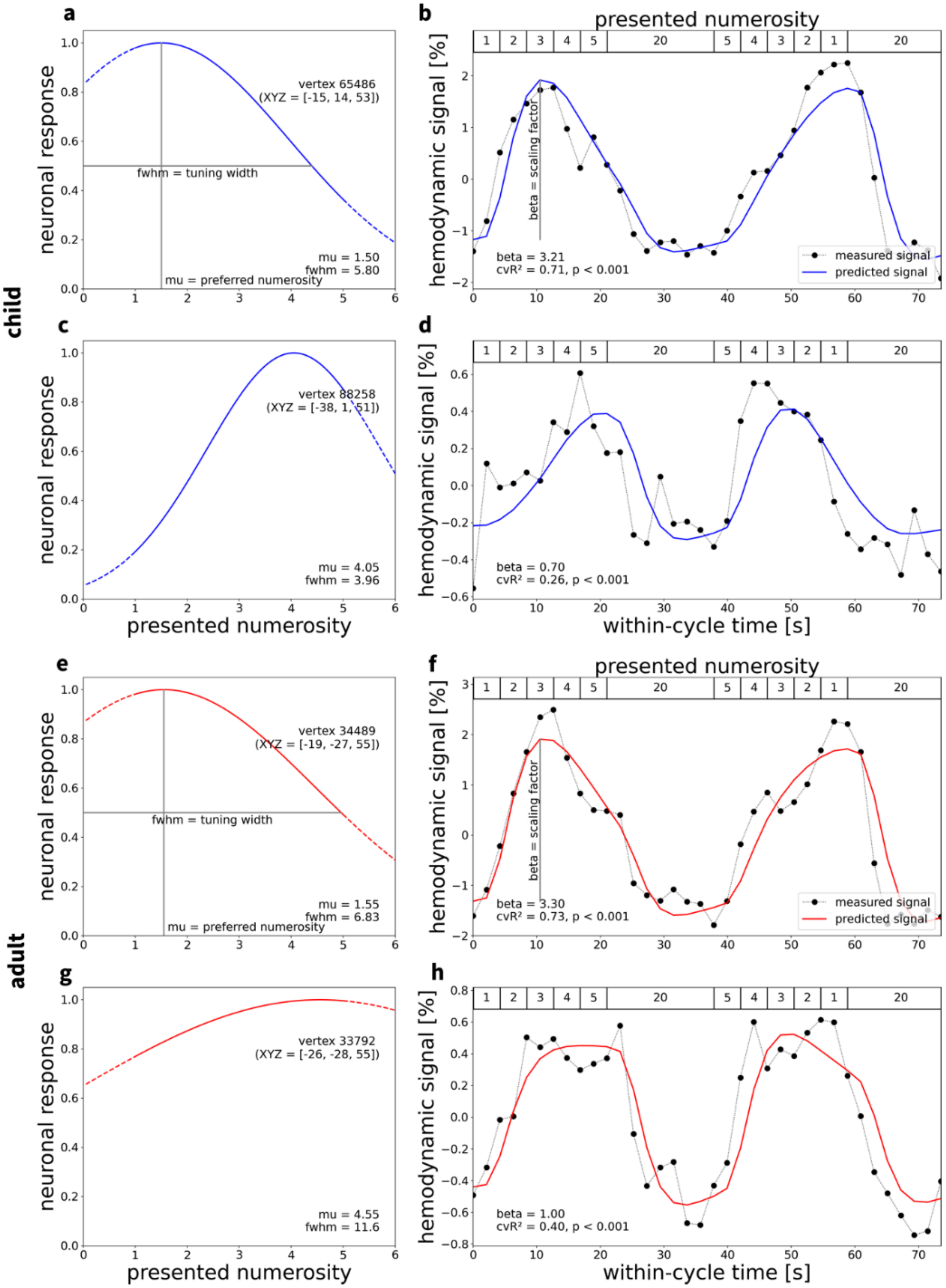


**Figure S9 | Linear tuning function results.** This figure follows Fig. 2a–h in terms of data and subjects, except that tuning functions describing hemodynamic responses were extracted using a linear model.

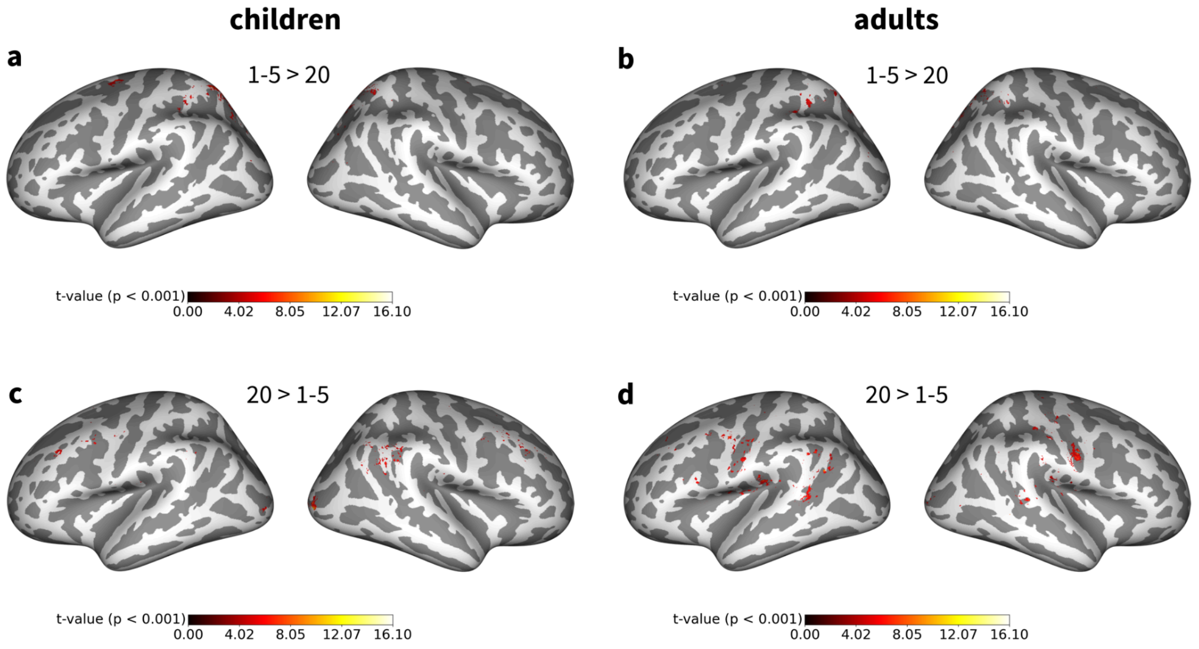


**Figure S10 | General Linear Model results.** Panels display thresholded statistical parametric maps ($p < 0.001$, uncorrected) corresponding to **a,b** significant positive effects of small numerosity (1-5 > 20) and **c,d** significant negative effects of small numerosity (20 > 1-5) for **a,c** children and **b,d** adults. The first colorbar tick label ($t = 4.02$) corresponds to the critical t-value for a one-sample t-test with sample size $n = 12$ and significance level $\alpha = 0.001$.

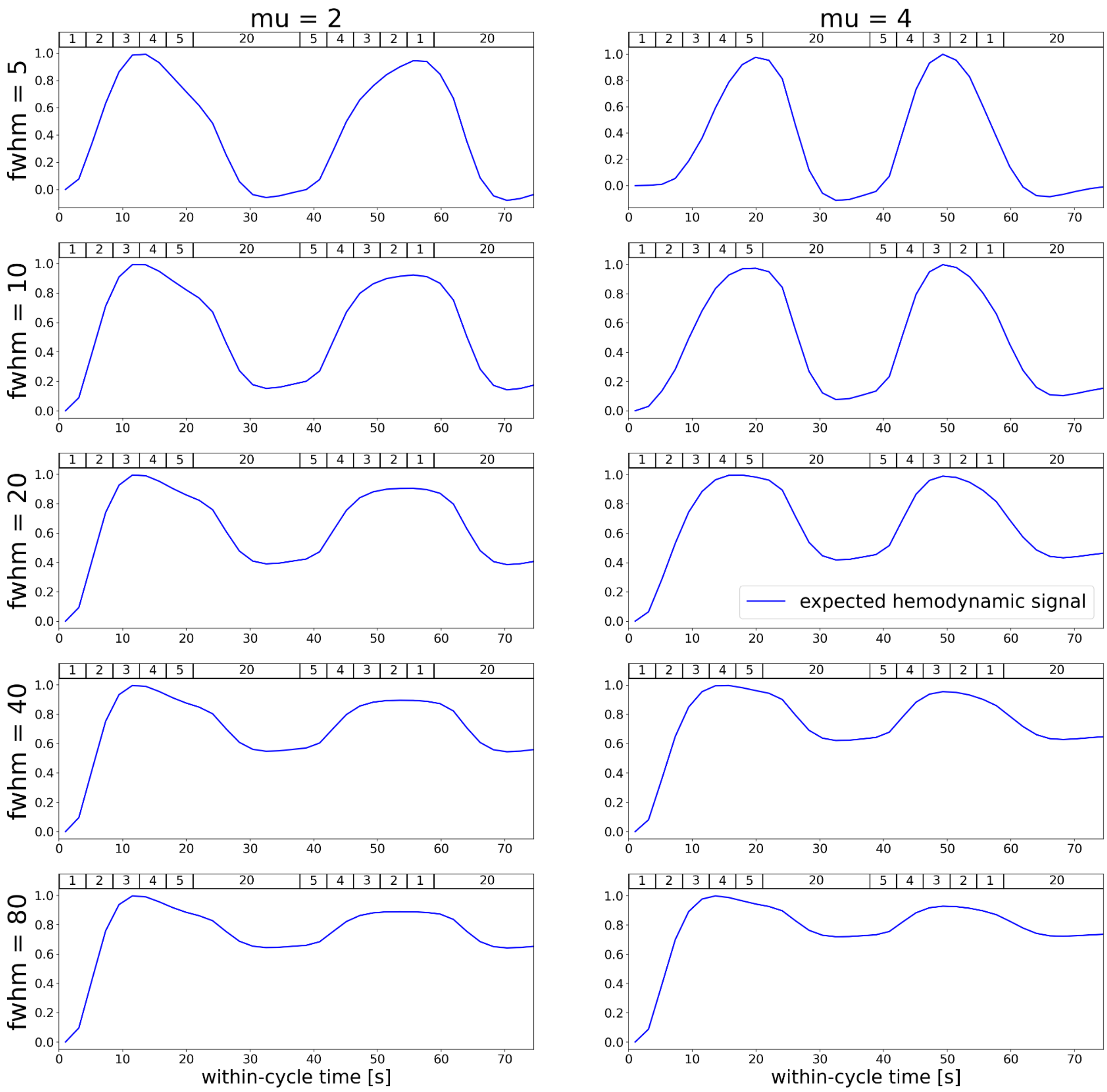


**Figure S11 | Simulated effect of tuning parameters on numerosity-related time courses.** Panels display expected hemodynamic time courses when generating neuronal signals given typical tuning parameters (logarithmic tuning function; left: mu = 2; right: mu = 4; from top to bottom: fwhm = 5, 10, 20, 40, 80; cf. Figure 5c,d) and convolving these signals with the canonical hemodynamic response function. These results show that differential responses to numerosities can emerge even with high tuning width for both, low and medium preferred numerosities.
